# Quantitative analysis of transcription start site selection in *Saccharomyces cerevisiae* reveals control by DNA sequence, RNA Polymerase II activity, and NTP levels

**DOI:** 10.1101/2021.11.09.467992

**Authors:** Yunye Zhu, Irina O. Vvedenskaya, Sing-Hoi Sze, Bryce E. Nickels, Craig D. Kaplan

## Abstract

Transcription start site (TSS) selection is a key step in gene expression and occurs at many promoter positions over a wide range of efficiencies. Here, we develop a massively parallel reporter assay to quantitatively dissect contributions of promoter sequence, NTP substrate levels, and RNA polymerase II (Pol II) activity to TSS selection by “promoter scanning” in *Saccharomyces cerevisiae* (Pol II MAssively Systematic Transcript End Readout, “Pol II MASTER”). Using Pol II MASTER, we measure the efficiency of Pol II initiation at 1,000,000 individual TSS sequences in a defined promoter context. Pol II MASTER confirms proposed critical qualities of *S. cerevisiae* TSS -8, -1, and +1 positions quantitatively in a controlled promoter context. Pol II MASTER extends quantitative analysis to surrounding sequences and determines that they tune initiation over a wide range of efficiencies. These results enabled the development of a predictive model for initiation efficiency based on sequence. We show that genetic perturbation of Pol II catalytic activity alters initiation efficiency mostly independently of TSS sequence, but selectively modulates preference for initiating nucleotide. Intriguingly, we find that Pol II initiation efficiency is directly sensitive to GTP levels at the first five transcript positions and to CTP and UTP levels at the second position genome wide. These results suggest individual NTP levels can have transcript-specific effects on initiation, representing a cryptic layer of potential regulation at the level of Pol II biochemical properties. The results establish Pol II MASTER as a method for quantitative dissection of transcription initiation in eukaryotes.

## Introduction

In transcription initiation, RNA polymerase II (Pol II), assisted by General Transcription Factors (GTFs), binds promoter DNA through interactions with core promoter elements. Subsequently, a turn of promoter DNA is unwound, forming a Pol II-promoter open complex containing a single-stranded “transcription bubble”, and Pol II selects a promoter position to serve as the transcription start site (TSS). At the majority of Pol II promoters in eukaryotes, TSS selection occurs at multiple positions ^1–11^. Thus, the overall rate of gene expression at the majority of Pol II promoters is determined by the efficiency of initiation from several distinct TSS positions. In addition, studies have suggested that alternative TSS selection can lead to differences in mRNA features, translation activity and subsequential protein levels and functions ^3, 12, 13^, and therefore is widespread in different cell types ^6^, developmental processes ^4, 12, 14, 15^, growth conditions ^16^, responses to environmental changes, and cancers ^17–19^.

Pol II initiation in yeast proceeds by a promoter scanning mechanism, where the Pol II Pre-initiation Complex (PIC), comprising Pol II and GTFs, assembles upstream of the initiation region and then scans downstream to select a TSS position ^20–27^. The efficiency of initiation at a given position depends on multiple features. Location relative to the core promoter region from which scanning will originate and proceed downstream, is critically important. DNA sequence in and around the TSS is critical for TSS specification. Here, the template base specifying the TSS position and the position immediately upstream of the TSS (positions +1 and -1, respectively) make the largest contributions. In particular, there is a strong preference for an R:Y base pair at position +1 and Y:R base pair at position -1 (reflected as a Y_-1_R_+1_ “initiator” sequence on the coding strand; Y=pYrimidine; R=puRine), and this preference is near universal for RNAPs^1, 10, 11, 16, 21, 22, 28–44^.

It has long been recognized that DNA sequences at positions beyond the -1/+1 also contribute to Pol II initiation efficiency. However, the sequence preferences at these positions are difficult to determine from genomic usage alone. This difficulty is due to differences in individual promoter contexts and may also include species-specific determinants. For example, in *S. cerevisiae*, genome-wide transcriptomic data have revealed a preference for an A:T base pair at position -8, reflected as an A on the transcribed strand and a T on the template strand ^1, 16, 21, 34, 35^. Furthermore, promoters with high expression and/or with initiation focused primarily at a single, efficient TSS tend to show additional A enrichment at positions -7 to -3 on the transcribed strand (T on the template strand) ^1, 16, 21, 45, 46^. Several factors confound efforts to directly measure the contribution of DNA sequence to the efficiency of initiation by Pol II, including promoter chromatin context and exact TSS position within a promoter. These attributes together might be considered a promoter’s architecture. Promoter architectural factors are especially apparent in yeast – where initiation occurs by promoter scanning – because TSSs are proposed to be selected in a polar fashion from upstream to downstream.

An elegant and important study of promoter scanning from Kuehner and Brow established that TSS usage is determined by TSS priority during the scanning process ^22^. As noted above, scanning proceeds directionally from upstream sequences to those downstream. Therefore, sequences are examined by the transcription machinery in the order in which they are scanned with upstream TSSs having priority, regardless of innate TSS strength. Kuehner and Brow introduced the concept of “TSS efficiency,” which accounts for how much Pol II reaches a particular TSS in order to compare innate TSS strengths between TSS sequences (see **Figure 1**). TSS efficiency is defined as the usage of a particular TSS normalized to the usage of that position and all downstream positions. Downstream TSS usage is used to estimate the flux of scanning Pol II that scans past a particular TSS. This measure is essential for comparison of TSSs embedded in different promoter contexts and different positions relative to other TSSs.

**Figure 1.**
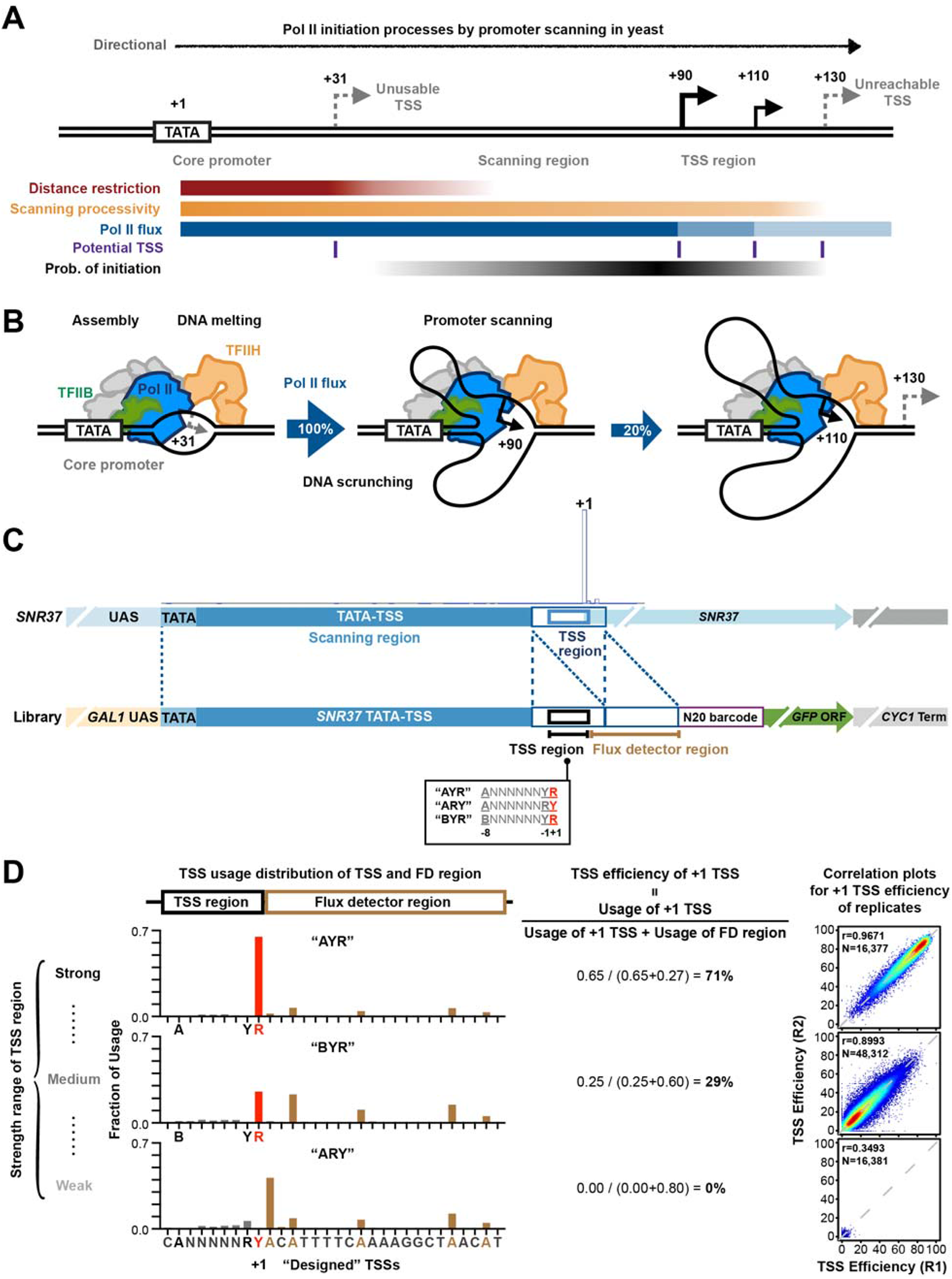
A high-throughput system for studying transcription TSS selection. **(A and B)** Pol II initiation in yeast proceeds by a promoter scanning mechanism. Yeast Pol II initiation usually occurs at multiple TSSs ranging from 40 to 120 bp downstream of a core promoter region comprising the PIC assembly position (the TATA-box for TATA-containing promoters). After the PIC assembles upstream (core promoter) scanning will proceed over a scanning region towards positions where TSS selection occurs (TSS region). In addition to sequence control of initiation through suitable TSS sequences, the probability of initiation across promoter positions is also controlled by multiple architectural features shown in **(A)**. These include the inhibition of initiation near a core promoter that diminishes as scanning proceeds (“distance restriction”), biochemical restrictions on how far scanning can proceed that increase as scanning proceeds (“scanning processivity”), and “Pol II flux”, which represents the decrease in amount of scanning Pol II as scanning proceeds due to conversion of scanning Pol II to transcribing Pol II upon initiation. **(C)** Construction of promoter libraries that control TSS sequence context. Top panel shows schematic of the *SNR37* promoter and its TSS usage distribution based on TSS-seq ^21^. *SNR37* has a focused and highly efficient TSS region. Bottom panel shows schematic of the Pol II MASTER libraries used in this study. A duplication of the *SNR37* TSS region was inserted before native TSS region, and the -8 to +1 positions relative to native *SNR37* +1 TSS (black box) were replaced by a 9 nt randomized region to provide almost all possible sequences. The second *SNR37* TSS region functions as a “Flux Detector” (FD) to capture Pol II flux that fails to initiate within the randomized region and allow determination of initiation efficiency within the randomized region. A barcode region (purple box) allows RNA products to be assigned to respective promoter variants. The *GAL1* UAS allows control of library expression, and the *GFP* ORF and *CYC1* terminator support termination and stabilize RNA products. **(D)** TSS usage distributions at TSS and FD regions for different promoter variant “AYR”, “BYR”, and “ARY” libraries are shown on the left. TSS usages from designed +1 TSS and positions upstream are in red and grey, respectively. TSS usage from the FD region is in brown. The definition of “TSS efficiency” and overall TSS efficiency calculations for the aggregate +1 TSS of all in different libraries are shown in the middle. Example correlation plots of TSS efficiency calculations for +1 TSSs from individual promoter variants in Pol II MASTER libraries between representative biological replicates are show on right. Pearson r and number of compared variants are shown.

Imbalanced promoter sequence distributions imposed by evolutionary processes also limit the ability to determine sequence-activity relationships for initiation. For example, it has been observed that yeast promoters have uneven distribution of bases across promoters and this is most obvious in the enrichment at highly expressed promoters for T on the transcribed strand upstream of the median TSS position (reflecting the middle of the TSS distribution) and A on the transcribed strand downstream of the median TSS position. Furthermore, yeast promoters have a paucity of G/C content in general ^21, 45–48^. Therefore, the biased distribution of bases in yeast promoters leads to a biased distribution of TSS sequence motifs. Preferred TSS motifs (those with a -8A) show enrichment downstream of promoter median TSS position, and less-preferred motifs (those without a -8A) show enrichment upstream of the median TSS position ^21^. We have found that hyperactive Pol II mutants (“gain-of-function” or GOF) and hypoactive Pol II mutants (“loss-of-function” or LOF) shift TSS usage distributions upstream or downstream, respectively, while also showing differences in aggregate usage of particular TSS motifs. Differences in usage cannot be disentangled from actual differences in TSS efficiencies due to biased promoter sequence distributions. Moreover, other properties such as the biochemistry of transcription itself due to availability of NTP substrates, the biochemical properties of scanning processivity ^20, 25, 27^, promoter identity ^49–51^, or promoter chromatin could also contribute to initiation output ^52^. Together, all of these factors would make contributions of primary DNA sequence difficult to ascertain in the absence of controlling for TSS context.

In order to remove contextual differences among promoters we have developed a system to dissect determinants of initiation efficiency within a defined, controlled promoter context. Here, we present “Pol II MASTER” based on bacterial MASTER (MAssively Systematic Transcript End Readout) ^42, 53–56^, which allows determination of initiation at base pair resolution and attribution of RNA transcripts to barcoded promoter DNA variants in massively parallel fashion. We apply Pol II MASTER to initiation by promoter scanning to investigate the initiation efficiency of ∼80,000 promoter variants within an appropriate TSS region in Pol II WT and catalytic mutants and upon manipulation of NTP levels. We show that this system enables determination of the interface between initiation factor activity, transcription substrates, promoter sequence, and promoter output. We recapitulate the previously defined impact of TSS positions -8, -1, and +1 on initiation efficiency while revealing the wide range of effects other positions have in initiation (-11 to -9, -7 to -2 and +2 to +4). We identify a distinct hierarchy in Y_-1_R_+1_ preferences, and detect interactions between bases at neighboring positions, suggesting potential mechanistic coupling between positions. We find that Pol II mutant classes increase or decrease initiation efficiencies for all possible sequences. This result is consistent with predictions that the most, but not all, effects of altered TSS selection in Pol II active site mutants (directional shifts in TSS distributions) are driven by initiation efficiency changes across all sequences and not on TSS sequence selection *per se*. Our results further show that Pol II activity level contributes to selective efficiency of initiation at sequences +1A vs +1G and find that initiation efficiency is selectively sensitive to individual NTP levels across genomic TSSs. Our findings demonstrate that Pol II MASTER provides a platform to quantitatively dissect how initiation factor activity together with promoter sequence contribute to Pol II transcription initiation in vivo.

## Results

### A high-throughput system for studying TSS sequence effects on Pol II initiation output

TSSs are identified within yeast promoters by scanning from upstream near the core-promoter to downstream (**Figure 1A**). While promoter melting occurs around +20 from the TATA box (if present), TSS usage is restricted adjacent to this region, with most TSS selection occuring ∼40-140 nt downstream from the core promoter ^57^ (**Figure 1B**). Processivity of the scanning process, likely determined by TFIIH activity, is expected to limit TSS usage at the downstream edge of promoters (**Figure 1A**, “unreachable TSS”). Once initiation happens at any site within this window of opportunity for initiation (**Figure 1A, B**), Pol II “flux” – the amount of Pol II progressing to TSSs downstream – is necessarily reduced. Given these potential variables in TSS context and the “first come-first served” basis of promoter scanning, determining how some TSSs are stronger than others cannot be accurately determined solely from TSS usage at genomic promoters. We have established a massively parallel promoter variant assay “Pol II MASTER” in order to dissect how TSS sequence controls initiation efficiency and how sequence interacts with Pol II catalytic activity or NTP levels in a controlled context. In Pol II MASTER, we have embedded almost all possible sequences within a 9 bp randomized TSS region (**Figure 1C**) constructed on plasmids and introduced these promoter libraries into yeast strains with wild type (WT) or mutated Pol II. The sequence libraries constructed are illustrated in **Figure 1C** and are referred to by their base compositions relative to the transcribed strand unless specifically noted otherwise. The libraries are named by identities of bases at positions -8, -1, and +1. The “AYR” library has composition A_-8_NNNNNNY_-1_R_+1_ (N=A, C, G, or T, Y=C or T, R=A or G) relative to the transcribed strand, with “BYR” having composition B_-8_NNNNNNY_-1_R_+1_ (B=C, G, or T), *etc*. Our three libraries comprise 81,920 promoter variants. Because each of these promoter variants were evaluated for TSS initiation efficiency at up to 12 positions within or adjacent to the randomized region, our assay allows for analysis of up to 983,040 distinct TSS sequences.

This randomized TSS region was inserted into a controlled promoter context containing specific functionalities (**Figure 1C**). <u>First</u>, the *GAL1* UAS was utilized to allow expression control of libraries. <u>Second</u>, the TATA-box-to-TSS region of *SNR37* gene was used as a “scanning region” to direct initiation within the downstream randomized TSS region. This is because almost no RNA 5′ ends are observable from this scanning region within its normal promoter context. <u>Third</u>, the native, highly-efficient *SNR37* TSS region was inserted downstream of the randomized TSS region as a “Flux Detector” (FD). Here, we employ the approach of Kuehner and Brow that a highly efficient initiation region placed downstream an upstream TSS region may capture any polymerases that happen to scan past the randomized TSS region. As a metric to compare strength of different TSSs, we employ “TSS efficiency” as defined by Kuehner and Brow ^22^, which measures the usage of a TSS relative to usage at that TSS and all downstream positions (**Figure 1C**). This metric serves two key purposes: it allows upstream and downstream starts to be compared as it takes priority effects into account (upstream TSSs reduce the amount of polymerases scanning to downstream sites), and it allows normalization of promoter usage using RNA only, engendering comparison across promoters and libraries. <u>Fourth</u>, a 24 bp DNA barcode containing 20 positions of randomized bases and 4 interspersed fixed bases (to exclude low-complexity sequences) allows RNA products to be assigned to respective promoter variants. An RNA barcode is critical as bases upstream of the TSS will not be present in the transcribed RNA yet still contribute to TSS efficiency. <u>Fifth</u>, the *GFP* coding region and *CYC1* terminator were added to support termination and stabilize RNA products. Libraries were constructed by PCR sewing followed by cloning into a plasmid backbone. After amplification in *E. coli*, plasmid libraries were transformed into Pol II WT and mutant yeast strains in triplicate. Library expression was induced by addition of galactose to the medium (4% final) for three hours. Both plasmid DNA and RNA products were extracted from harvested yeast cells and amplified for DNA-seq and TSS-seq (**Figure S1A**).

Several measures indicate high level of reproducibility and coverage depth of library variants (**Figure 1D, Figure S1**). Base coverage in the randomized region was highly even (**Figure S1B**). Correlation analysis for DNA-seq reads per promoter variant suggested that yeast transformation did not alter promoter variant distribution (**Figure S1C, D**). Bulk primer extension of libraries illustrated their average behavior and the amount of initiation deriving from the randomized region (**Figure S1E**). As designed, only a very small fraction of initiation was generated from the barcode region or further downstream, validating that the flux detector captured scanning polymerases (**Figure S1E, F**). Aggregate distribution of reads in our three libraries shows that as TSSs decreased in efficiency from the most efficient library (“AYR”) to least (“ARY”) reads shift from the designed +1 TSS to downstream positions (**Figure 1D**, left). The apparent shift of TSS usage to position -1 in the ARY library was because the purine at the designed -1 position serves as a +1 for newly created TSSs. **Figure 1D** (middle) illustrates the aggregate TSS efficiency of each library based on usage at the designed +1 TSS relative to usage at that position plus all downstream usage. Correlation analysis for efficiency of library “major” TSSs (designed +1 TSS of “AYR” and “BYR” libraries, +2 TSS of “ARY” library) demonstrated that biological replicates were highly reproducible (**Figure 1D**, right). Therefore, we summed reads from the three biological replicates, keeping TSSs that contained at least five TSS-seq reads in each replicate and whose Coefficient of Variation (CV) in TSS-seq reads across replicates was less than 0.5 (as a proxy for reproducible behavior) (**Figure S1G**). As a result, 97% of possible TSS promoter variants were covered in each library on average (**Table S3**). Finally, we analyzed potential interactions between TSS sequences and overall promoter expression (**Figure S1H**). Normalization of individual promoter output (RNA levels) to promoter template number (DNA level) indicates that total promoter output based on TSS usage across each promoter was relatively unaffected by individual TSS strength within our randomized region.

### Sequence-dependent control of TSS efficiency in *S. cerevisiae*

To ask how our libraries recapitulated known TSS efficiency measurements, we first examined core sequences in our library matching the *SNR14* TSS and its variants previously analyzed by Kuehner and Brow ^22^ for TSS efficiency (**Figure 2A**). Our randomized library contains the *SNR14* TSS sequence embedded in our *SNR37* context along with all single substitution variants of this sequence, including the subset previously examined at *SNR14.* We found that Pol II MASTER recapitulated the single base effects on TSS efficiency previously observed while also indicating that single base changes around a TSS can have large effects on TSS efficiency.

**Figure 2.**
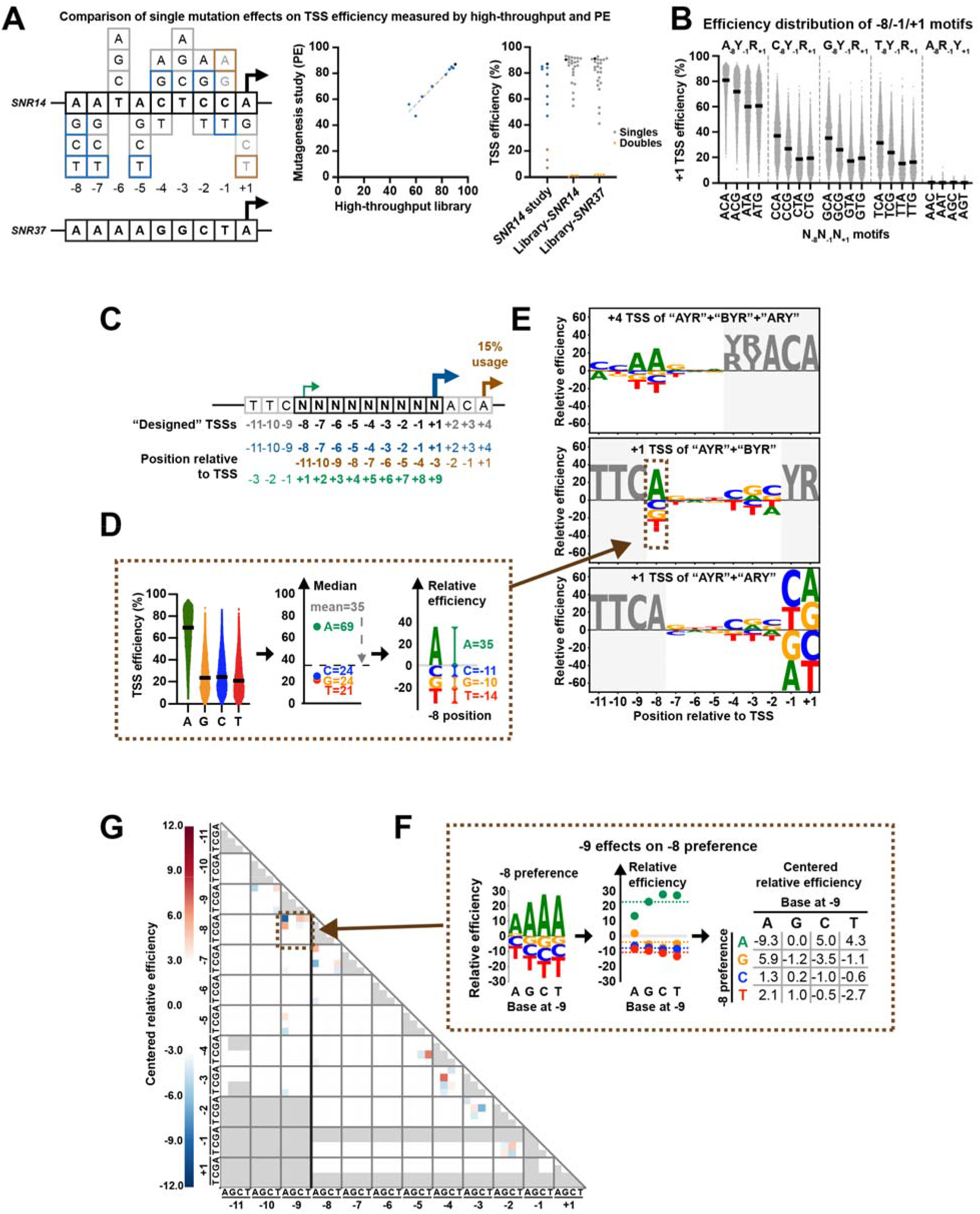
TSS and surrounding sequences modulate initiation efficiency over a wide range. **(A)** Comparison of single base mutation effects on TSS efficiency measured by Pol II MASTER and primer extension. (Left) Sequences of *SNR14* and *SNR37* TSS regions (in black boxes, including positions between -8 to +1 relative to TSS) and all possible single substitutions of *SNR14* TSS region. Single substitutions included in both a prior *SNR14* mutagenesis study ^22^ and Pol II MASTER libraries are in blue while those lacking in our study are in brown. Additional substitutions analyzed here are in gray. (Middle) High correlation of TSS efficiency measured by Pol II MASTER and primer extension. Mutation classes color coded as on left. (Right) Range of effects of single base substitutions on TSS efficiency for *SNR14*- and *SNR37*-related sequences. Mutation classes color coded as on left. Single substitutions absent from Pol II MASTER because of library design (R_-1_ and Y_+1_) were expected to have super low TSS efficiencies. Double substitutions of *SNR14* and *SNR37* TSS region included in Pol II MASTER “ARY” library are shown as orange inverted triangles and show almost no efficiency. **(B)** Pol II initiation shows a strong preference for A_-8_ and C_-1_A_+1_ containing variants. All promoter variants were divided into 20 groups defined by bases at positions -8, -1 and +1 relative to the designed +1 TSS, and their +1 TSS efficiencies were plotted as spots. Lines represent median efficiencies of each group. **(C)** Definition of “designed” +1 TSS (designated as +1) and positions relative to this TSS (blue TSS arrow and sequence). TSS usage generated from upstream or downstream of “designed” +1 TSS (green and brown TSS arrows and sequences, respectively) allows us to study sequence preferences at positions -11 to -9 and +2 to +9 relative to site of initiation. **(D)** Schematic illustrating how “relative efficiency” is calculated and visualized as a sequence logo in **E**. At a particular position relative to a particular TSS, first, all variants were divided into the four base subgroups defined by base at this position. Next, median values of each base group were extracted and centered based on the mean of all median values. The centered median values were defined as “relative efficiencies”, representing preferences for bases at this particular position. Finally, relative efficiencies of bases were visualized as sequence logos. Positive and negative values indicate relatively preferred or less preferred bases, respectively. **(E)** Pol II initiation shows distinct sequence preference at positions around the TSS. Top panel was generated using datasets of designed +4 TSS deriving from “AYR”, “BYR” and “ARY” libraries. Middle panel was generated using datasets of designed +1 TSS deriving from “AYR” and “BYR” libraries. Bottom panel was generated using datasets of designed +1 TSS deriving from “AYR” and “ARY” libraries. Positions that contain fixed or not completely randomized bases are shown in grey. **(F)** Schematic illustrating how sequence interaction between positions is calculated and visualized as a heat map in **G**. Using -9/-8 positions as an example, relative efficiencies at position -8 were calculated when different bases were present at position -9. Next, relative efficiencies of each base were centered based on the mean of all relative efficiencies of a particular base. After centralization, negative and positive values indicate negative and positive interactions. Interaction scores for two sequence positions are read at the intersection of the *x* and *y* axes labeled by base and position. That means, for example, -9.3 at -9A/-8A and 5.9 at -9A/-8G intersections indicate having a -9A decreases A preference and increases G preference at position -8, respectively. Finally, the centered relative efficiencies matrix was visualized as a heat map to represent the interaction between examined positions. **(G)** Sequence interactions are mainly observed at neighboring positions. Red and blue indicate positive and negative interactions, respectively. Missing values are shown in grey. Interactions related to positions -11 to -9 were calculated using datasets of designed +4 TSS deriving from “AYR”, “BYR” and “ARY” libraries. Other interactions were calculated using datasets of designed +1 TSS deriving from “AYR” and “BYR” libraries.

Examining all designed +1 TSS variants present in our library, we first focused our analysis on positions -8, -1 and +1, which *in vivo* genome-wide data suggested are important determinants for TSS selection (**Figure 2B, Figure S2A**). We determined the TSS efficiencies of the designed +1 TSS for all promoter variants, dividing all variants into 64 groups defined by bases at positions -8, -1 and +1 relative to TSS. This allowed us to examine the TSS efficiency distribution within each subgroup. Our results demonstrated the known importance of these three positions but within our defined promoter context, but also demonstrated that surrounding positions have a considerable impact on TSS efficiency (note the wide range of efficiencies within each TSS group, **Figure 2B**). <u>First</u>, in our controlled context for TSSs at the designed +1 position, we found that Y_-1_R_+1_ was essential for initiation above a minimal background relative to R_-1_Y_+1_. Even in the presence of an A_-8_, R_-1_Y_+1_ variants showed essentially no usage. <u>Second</u>, we quantified the very large effect of a -8A on TSS efficiency (note that A_-8_Y_-1_R_+1_ motifs were much higher in efficiency in aggregate than non-A_-8_Y_-1_R_+1_ TSSs), clearly demonstrating that -8A alters TSS efficiency. This result is in line with the observation that A_-8_C_-1_A_+1_ motif-containing TSSs have the highest aggregate TSS usage from genomic promoters and appear to be the most efficient ^21^. <u>Third</u>, among Y_-1_R_+1_ elements we found a clear hierarchy of efficiency – C_-1_A_+1_ > C_-1_G_+1_ > T_-1_A_+1_ ≈ T_-1_G_+1_ – that could not be discerned from genomic promoter TSS usage or contexts, likely due to promoter sequence biases within genomic promoters.

As noted above, a wide range of TSS efficiencies were observed within every -8/-1/+1 group, suggesting contributions of other sequence positions to initiation efficiency. To determine if base identities within -7 to -2 positions have similar effects regardless of -8, -1 and +1 identity, we rank ordered all individual TSSs for each N_-8_N_-1_N_+1_ motif by the efficiency of their A_-8_C_-1_A_+1_ version (**Figure S2A**). This rank ordering by A_-8_C_-1_A_+1_ efficiency was predictive of efficiency ranks for -7 to -2 sequences with different bases at positions -8, -1 and +1. This observation indicates important contributions from positions beyond positions -8, -1 and +1 and that positions might function independently to determine TSS efficiency (examined below).

We set out to determine the contributions to Pol II TSS efficiency of individual bases at each position across our randomized region relative to the designed +1 TSS. To do so, we calculated the TSS efficiencies of examined TSS variants and divided them into subgroups based on bases at individual positions relative to the designed TSS +1 position (**Figure S2B**). Comparison of TSS efficiencies across base subgroups suggested significant individual base effects on TSS efficiency in aggregate at all examined positions. The usage of other positions in our promoter outside of the designed TSS +1 position (either within the randomized region or outside of it) allowed us to examine contribution of bases in our randomized region to the efficiency of these other TSSs (**Figure S1E, F,** **Figure 2C**). Our combined libraries allowed us to analyze efficiencies of nearly a million TSS variants present within them (distribution of efficiencies is shown in **Figure S2C**). In order to visualize sequence preferences, we used the median initiation efficiency values of each base subgroup as indicators for preference, using centered median values to calculate “relative efficiency” and illustrated this preference in a sequence logo (**Figure 2D**). Datasets of designed +1 TSSs deriving from “AYR”, “BYR”, and “ARY” libraries allowed us to nearly comprehensively study preference at positions -8 to +1 (**Figure 2E**, middle and bottom). Additionally, 10-25% of total TSS usage among different libraries deriving from a TSS at +4 (**Figure 2C**, brown TSS arrow and sequences) allowed study of positions -11 to -9 relative to this TSS (**Figure 2E**, top). As noted above, positions -8, -1 and +1 were major determinants for TSS efficiency. Interestingly, position -9 showed a relatively strong effect in our defined promoter context, which was not obvious from genome-wide analyses. At positions -4 to -2, we observed modulation of initiation efficiency, where in general C and/or G were preferred and T was less-preferred. This overall preference is consistent with an observation where individually mutating Ts at positions -4 to -2 relative to -38 TSS of *ADH1* promoter to a C significantly increased usage of that TSS ^57^. Though preferences at positions -7 to -5 were statistically significant (**Figure S2B**), contributions were much lower than other examined positions. Taken together, these results indicate that positions -9 to -8 and -4 to +1 are two major sequence position groups contributing to TSS efficiency.

Experiments across species and the description of the canonical initiator element (Inr) suggest some sequence contributions from downstream positions relative to the TSS ^10, 43, 44, 57–61^. To determine the impact of sequences downstream of the TSS, we examined motif enrichment of the most efficient -8 TSS variants, whose positions +1 to +9 are located in the randomized region (**Figure 2C**, green TSS arrow and sequences). We found an A(/G)_+2_G(/C)_+3_G(/C)_+4_ enrichment for the top 10% most efficient TSS, but not for the next 10% most efficient TSS (**Figure S2D**). These preferences are consistent with early mutagenesis work in yeast, where A(+2) to C/T, G(+3) to T, or C(+4) to T substitutions decreased the utilization of a particular TSS, but A(+2) to G or T(+5) to C substitutions had minor effects ^57^.

Pairwise nucleotide-position dependencies have been observed for some processes, for example in 5′ splice sites ^62–64^. To investigate potential higher order sequence interactions *i.e.* coupling between positions contributing to TSS efficiency, we examined all possible pairwise interactions among positions -11 to +1 (**Figure 2F, G, Figure S2E, F**). “Coupling” would entail a base at one position determining the contribution or effect of a base at another position. We found evidence for coupling at multiple positions with the strongest coupling between the -9 and -8 positions. Here, an A at one position suppresses the preference of A at the other position. This observation is indicative of positional epistasis at each position where an A at one position might diminish the impact of an A at the other (**Figure S2E**). Using this -9/-8 interaction as an example, **Figure 2F** shows how coupling was detected and visualized. We calculated “centered relative efficiency” values for each base and position and visualized them as a heatmap (**Figure 2F, G**). We found that evident interactions were mainly observed at neighboring positions (**Figure 2G**), especially within positions -9 to -7 and within positions -5 to -2. In addition to its interaction with position -9 described above, position -8 was also observed to interact with position -7 (**Figure S2F**). These observed interactions suggest that these positions might work together in TSS selection (see Discussion).

### Pol II mutants alter TSS efficiency for all possible TSS motifs while showing selective effects at +1

Pol II mutants were observed previously to change apparent specificity for A_-8_ versus B_-8_ (non-A) TSSs in opposite directions depending on Pol II defect by analysis of genomic TSS usage ^21^. As we noted in our prior work, this result could be a consequence of Pol II GOF or LOF mutants shifting TSSs upstream or downstream, respectively, coupled with the uneven distributions of bases and TSS motifs within genomic promoters. Furthermore, TSS motifs with a -8A by definition have an increased probability of having a potential upstream TSS that can be used *e.g.* the -8A could function as an upstream +1R TSS but might lack its own -8A. As a result, upstream-shifting mutants could *seem* to decrease preference for -8A TSSs because they are shifting from these to use of the -8A as a +1A. To determine how Pol II catalytic activity affects TSS selection in our controlled promoter context, we measured effects on TSS efficiency between WT and Pol II mutants in our promoter variant libraries (**Figure 3, Figure S3**). We introduced variant libraries into two Pol II LOF mutants (*rpb1* F1086S and H1085Q) and two GOF mutants (*rpb1* E1103G and G1097D). Data for all mutants showed high coverage of promoter libraries (**Figure S3A**), were highly reproducible (**Figure S3B-E**), and showed expected upstream or downstream shifts in TSS usage in aggregate across libraries (**Figure S3B**) with broad effects on efficiencies across sequences (**Figure 3, Figure S3F-I**). In our previous work ^21^ we proposed that Pol II mutants with decreased catalytic activity (LOF) would decrease initiation across all sequences; conversely, alleles with increased catalytic activity (GOF) would increase initiation across all sequences. We find that this is indeed the case (**Figure 3A**). In our controlled promoter context, we find that LOF mutants decreased efficiencies across all sequences, and GOF mutants increased efficiencies across all sequences, and that mutants did not show strong effects on -8 in contrast to apparent effects based solely on TSS usage from genomic promoters ^21^. Instead, we observed differences in Pol II mutants for efficiency of +1A TSS sequences relative to +1G TSSs (**Figure 3A**). These effects were apparent across the range of TSS efficiency (**Figure 3B**). Both GOF and LOF mutants showed stronger effects on +1A relative to +1G, consistent with the potential for changes to the Pol II active site to alter activity for different initiating NTPs.

**Figure 3.**
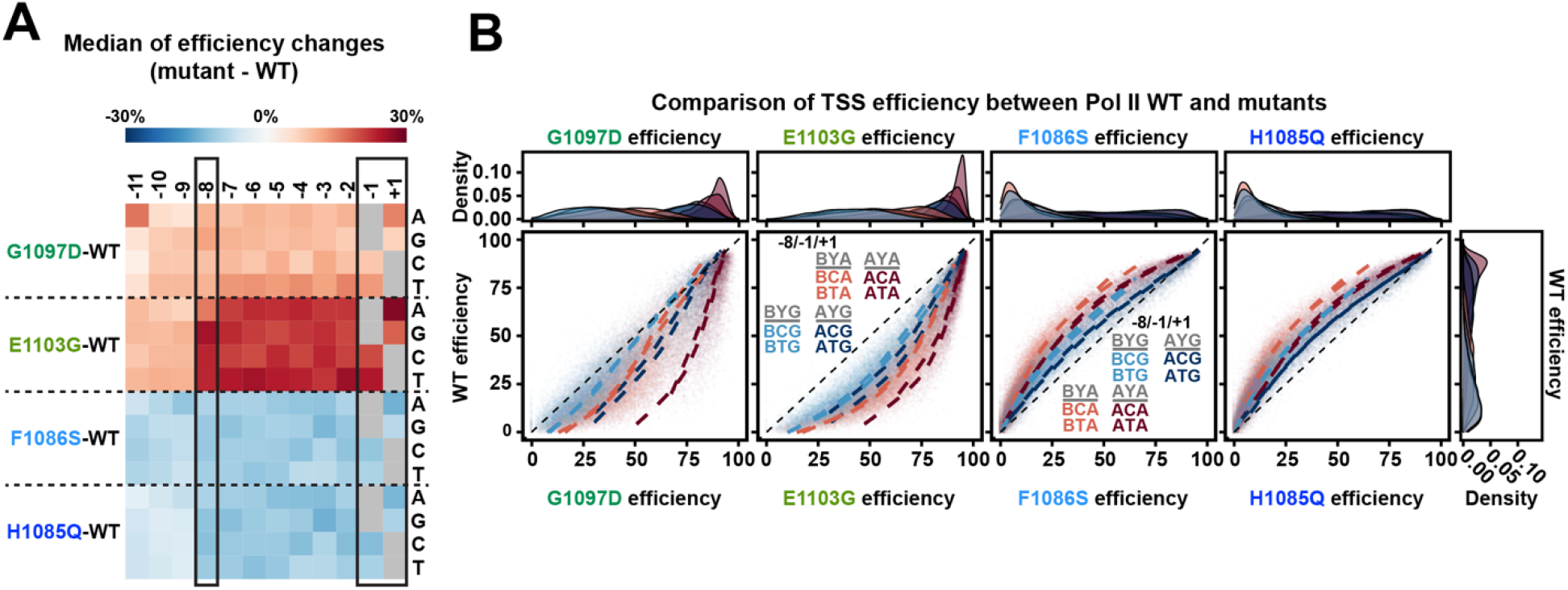
Pol II mutants alter TSS efficiency for all possible TSS motifs while showing selective effects for base at +1. **(A)** Pol II mutants alter TSS efficiencies across all motifs, corresponding to their direction of change to Pol II catalytic activity *in vitro*. TSS efficiency changes for each TSS variant were first determined by subtracting WT efficiency from Pol II mutant efficiency. The medians of efficiency changes for variant groups with indicated bases at each position relative to TSS were then calculated and illustrated in a heat map. Positive (red) values indicate Pol II mutants increased overall efficiency while negative (blue) values indicate decreased overall efficiency. **(B)** WT TSS efficiency for TSS variants divided into motif groups are plotted for mutant TSS efficiencies for the same TSS groups. The eight possible groups of TSSs for A/B_-8_C/T_-1_A/G_+1_ motifs were plotted and curve fit. Histograms show density of variants within each -8/-1/+1 subgroups. As to position -8, A_-8_ containing motifs show higher efficiency than B_-8_ containing motifs in both Pol II GOF (G1097D and E1103G) and LOF (F1086S and H1085Q) mutants (A_-8_ motifs: maroon and blue vs B_-8_ motifs: light coral and light blue). This is consistent with the proposed function of -8A to retain TSSs longer in the Pol II active site during scanning. This means that -8A may boost the positive effects of GOF mutants, therefore Pol II GOF mutants showed greater effects on A_-8_ motifs compared to B_-8_ motifs. In contrast, -8A compensates for active site defects of LOF mutants, therefore Pol II LOF mutants showed reduced effects on A_-8_ motifs compared to B_-8_ motifs. Both GOF and LOF mutants show reduced effects on G_+1_ motifs relative to A_+1_ motifs (G_+1_ motifs: light coral and maroon vs A_+1_ motifs: light blue and blue).

### Pol II initiation for +1G TSSs is sensitive to changes in GTP levels

We hypothesize that Pol II mutant alteration to the Pol II active site could result in altered substrate interactions. Therefore, such active site changes could result base-selective effects on TSS initiation efficiency with one +1-specified NTP, e.g. ATP, relative to another, e.g. GTP.. However, it is also conceivable that NTP levels within cells might be altered in Pol II mutants. To investigate if the observed selective effects of Pol II mutants for initiating NTP could be due to altered NTP levels, we measured total NTP levels in Pol II WT, F1086S (LOF), and E1103G (GOF) strains (**Figure 4A**). E1103G showed similar NTP levels as WT while F1086S increased NTP levels to different extents. While both ATP and GTP in F1086S were higher than WT, we reasoned that initiation might be more sensitive to GTP levels as GTP is less in excess of the apparent K_D_ of Pol II for NTPs than is ATP. If so, this could provide an explanation for the relatively smaller F1086S defect for +1G TSSs than for +1A TSSs; the increased GTP level in F1086S may partially buffer F1086S decreases in efficiency for +1G TSSs.

**Figure 4.**
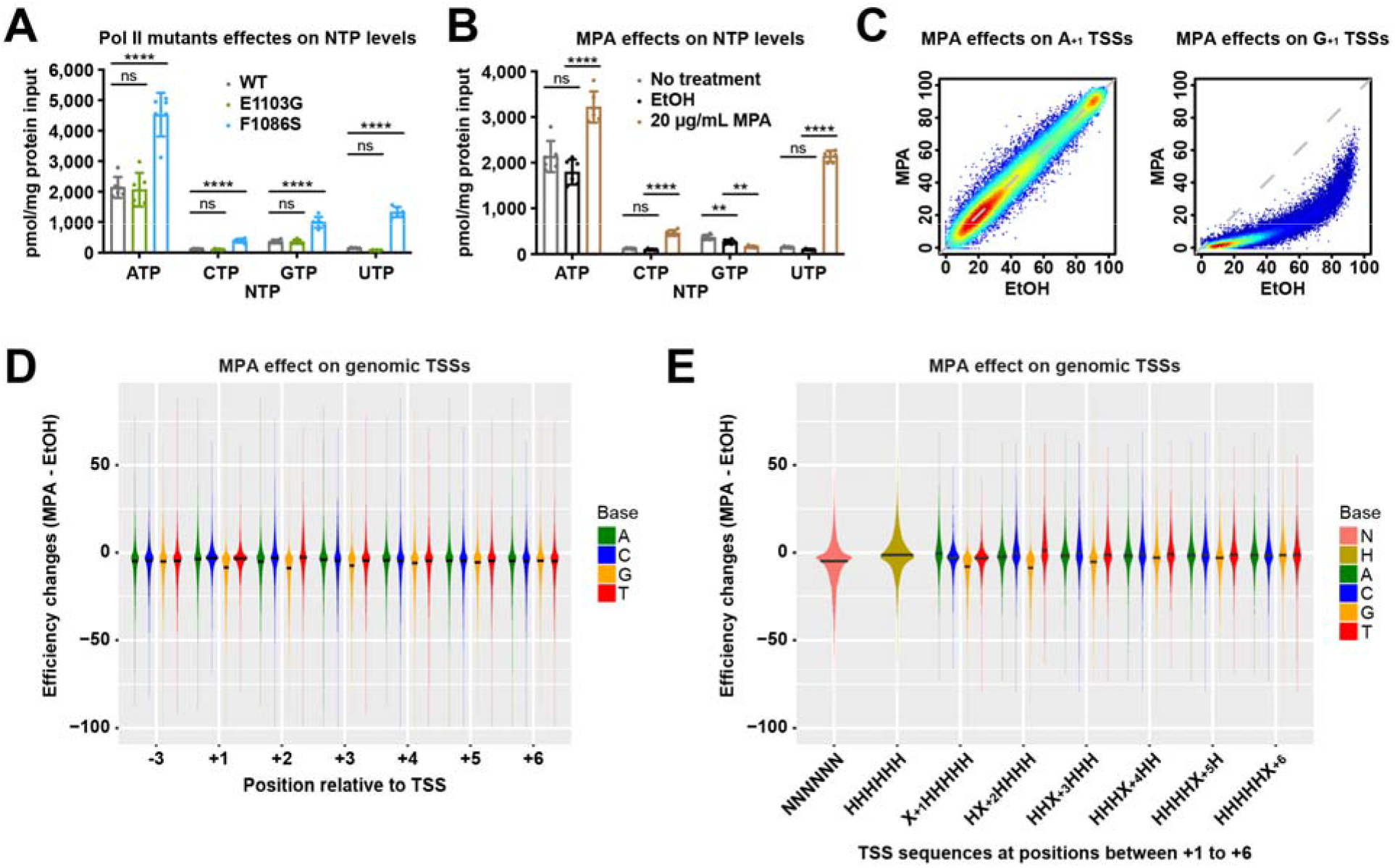
Pol II initiation is sensitive to NTP pools. **(A)** NTP levels measured in WT, Pol II E1103G (GOF) and Pol II F1086S (LOF) mutants. Statistical analyses by Ordinary one-way ANOVA with Dunnett’s multiple comparisons test for each NTP leave are shown. ****, P ≤ 0.0001; ***, P ≤ 0.001; **, P ≤ 0.01; *, P ≤ 0.05. **(B)** NTP levels measured in WT, WT treated with 100% ethanol (the solvent for MPA) and WT treated with 20 ug/ml MPA. Statistical analyses by RM one-way ANOVA with Dunnett’s multiple comparisons test for each NTP leave are shown. ****, P ≤ 0.0001; ***, P ≤ 0.001; **, P ≤ 0.01; *, P ≤ 0.05. **(C)** MPA selectively decreases TSS efficiencies of +1G TSSs (right) but shows no effects on +1A TSSs (left) in Pol II MASTER libraries. **(D)** MPA alters TSS efficiencies at genomic TSS positions. TSS efficiency change shown as difference between efficiency in EtOH and MPA treatment for all TSSs ≥ 2% efficiency in EtOH conditions within the 25%-75% of TSS distributions (see Methods). Bars on violin plots indicate medians computed based on density estimates. (**E**) Comparison of TSSs of any composition for first six nucleotides (NNNNNN) with those that lack G (HHHHHH, H=not G) or subsets of TSSs with A,C,G,T at positions +1 to +6. Bars on violin plots are as in (D). All C,G,T datasets for all positions 1-5 are distinct from A as a baseline comparison (Wilcoxon rank sum test with continuity correction). Position 6 each base is not significantly different from A.

If initiation were sensitive to GTP concentration in vivo, we hypothesized that reduction in GTP should result in selective reduction in initiation efficiency for +1G TSSs relative to +1A TSSs. Therefore, we examined if and how initiation of our promoter variant libraries and genomic TSSs was affected by treatment of cells with Mycophenolic Acid (MPA), which is known to deplete GTP through inhibition of yeast inosine monophosphate dehydrogenase (IMPDH) activities encoded by *IMD3* and *IMD4* ^65, 66^. We treated our mixed AYR and BYR libraries with MPA for 20 min prior to Gal induction, while EtOH treatment alone (the solvent for MPA) was used as a control. MPA decreased GTP level as expected while increasing ATP, CTP, and UTP levels (**Figure 4B**).

MPA showed effects on promoter variant libraries in aggregate (**Figure S4A**) with an overall downstream shift in TSS usage relative to EtOH alone. Comparison of replicates indicated high reproducibility of MPA treatment (**Figure S4A-B**). When examining specific sequences, we observed a striking and selective decrease in efficiency for essentially all +1G TSSs, while +1A TSSs were almost entirely unaffected (**Figure 4C**). MPA was previously shown to depress efficiency of +1G TSSs at *IMD2* as part of its regulation by GTP levels ^67^. Our results extend this observation to essentially all +1G TSSs in our MASTER library.

### Pol II initiation efficiencies are altered genome-wide upon changes to NTP levels

To determine if these striking effects were observed in promoter contexts beyond our designed libraries, we performed TSS-seq on genome-derived RNAs from the same samples that were analyzed for MASTER libraries. Examination of genomic TSSs revealed effects of MPA on TSS efficiencies across the genome (**Figure 4D**). We observed a depression of efficiency on average across all TSSs in MPA vs. EtOH (vehicle) treatment. Given that we observed strong effects at +1G for MASTER libraries, we speculated that there might also be effects on TSSs with +2G or potentially beyond, especially because initial bond formation requires interaction with two substrates, and subsequent initial elongation will lack stabilization of the product RNA that occurs with a full-length (∼9 nt) RNA:DNA hybrid. Examination of positions +2 through +6 indicates that TSSs with a G through position +5 are correlated with decreased initiation efficiency (**Figure 4D, S4D**). Conversely, we observed that presence of a +2C or +2U correlated with increased efficiency relative to +2A or +2G. These results are consistent with MPA-induced CTP and UTP increases promoting initiation efficiency for TSSs specifying C or U at +2. We hypothesized that A positions showed decreased overall efficiency because most transcripts might have at least one G within the first 5-6 nucleotides. To examine TSSs more carefully for G effects, we examined TSSs lacking G entirely for the first 6 positions (**Figure 4E**). TSSs lacking a G in the first 6 positions were more efficient on average than all TSSs in the presence of MPA explaining some of the overall suppression of all TSS efficiencies by MPA. This analysis also allowed us to demonstrate effects of a single G on initiation efficiency between positions +1 to +5..

### Learned initiation preferences are predictive of TSS efficiencies at genomic promoters

To ask how sequence determinants identified here relate to natural promoters, we compared our library-defined sequence efficiencies to TSS efficiencies observed at genomic promoters (**Figure 5**). To limit potentially confounding factors for genomic promoters, we focused on a single “median” TSS for each promoter in a defined set of promoter windows. The median TSS is defined as the TSS representing the position of 50^th^ percentile of reads within each promoter window ^21^, *i.e.* the TSS representing the middle of the cumulative distribution function of promoter reads from upstream to downstream. We found that Pol II sequence preference at positions around median TSSs was mostly consistent with what we observed in our libraries (**Figure 2E**, **Figure 5A**). Efficient genomic TSSs appear enriched for A at positions -7 to -5. The A-richness at positions between -10 to -3 and +5 to +10 has been noted in previous studies from our lab ^21^ (**Figure 5B**) and others ^1, 16, 34^. However, As at positions -7 to -5 appeared neutral in our promoter libraries (**Figure 2E**). The observed A-richness in the genome could reflect selection *in vivo* for additional promoter properties, such as providing an easily meltable DNA region, lower nucleosome occupancy or reflect a context dependent role not reflected by our promoter libraries.

**Figure 5.**
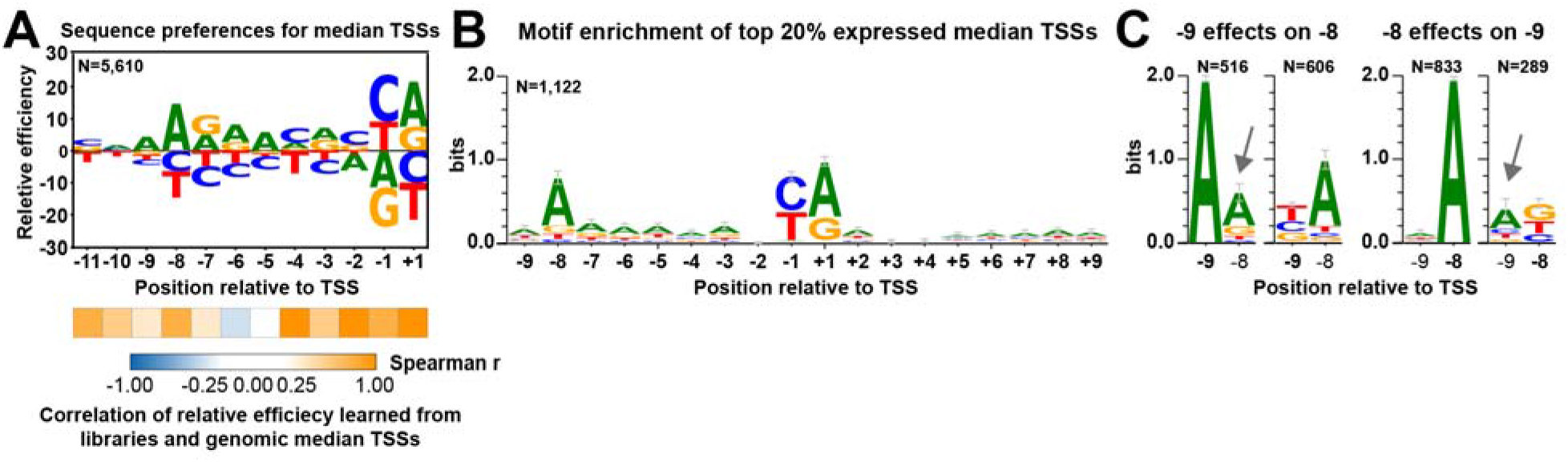
Learned initiation preferences are predictive of TSS efficiencies at genomic promoters. **(A)** Sequence preferences determined from TSSs representing the median of the distribution of usage from a set of 5979 yeast promoters (“median TSSs”) are congruent with library determined TSS preferences, except there is preference for A at positions -7 to -5 learned from genomic data. Sequence context and TSS efficiency of median TSSs were extracted from genomic TSS-seq data (GSE182792) ^27^. Calculation and visualization were as performed for promoter variant libraries. The number (N) of genomic median TSSs examined is shown. Statistical analyses by Spearman’s rank correlation test between relative efficiency at individual positions learned from promoter variant libraries and genomic median TSSs are shown beneath the sequence logo. **(B)** A-richness upstream and downstream of TSS is observed for highly expressed median TSSs. The number (N) of analyzed median TSSs is shown. Bars represent an approximate Bayesian 95% confidence interval. **(C)** Having an A at either position -9 or -8 reduces enrichment of A at the other position. Top 20% expressed median TSSs were divided into subgroups based on bases at position -9 or -8. Motif enrichment analysis was individually performed to subgroups. Numbers (N) of median TSSs within each subgroup are shown. Bars represent an approximate Bayesian 95% confidence interval.

We find that the interaction between -9A and -8A discovered in our libraries is also reflected in genomic TSSs (**Figure 5C**). We first grouped the median TSSs from the top 20% expressed promoters based on the base at position -9 or -8 and then examined sequence enrichment at the other position. We observed that -9A decreased the enrichment of -8A relative to that when position -9 is not A (**Figure 5C**, left). Moreover, when A was absent from the position -8, a higher enrichment for -9A was observed (**Figure 5C**, right). These results suggest that -9A may function in similar fashion as -8A, but that -8A may be more effective and therefore has been evolutionarily favored (see Discussion).

### Regression modeling identifies key DNA sequences and interaction for TSS selection regulation

We have found that DNA sequences around the TSS not only additively but also interactively contribute to TSS efficiency. To quantitatively identify key features (sequences and interactions) for TSS efficiency, we compiled datasets deriving from all libraries and predicted TSS efficiency from sequence information by logistic regression coupled with a forward stepwise selection strategy (**Figure 6A-E, Figure S5A-C**). We first compiled datasets generated from designed -8 to +2 and +4 TSSs deriving from “AYR”, “BYR”, and “ARY” libraries (**Figure S5A**) and split the data into training (80%) and test (20%) sets. For each variant, its sequences at positions -11 to +9 were extracted as potential predictors for TSS efficiency. We then used a forward stepwise strategy with a 5-fold Cross-Validation (CV) to select robust features (predictors). By evaluating model performance with R^2^, sequences at nine positions (positions -9 to -7 and -4 to +2) and one interaction (-9/-8 interaction) were identified as robust features and selected for final modeling (**Figure 6A**). It is worth noting that models with as few as three features – sequences at positions -9, -8, -1 (or +1) – could explain 74.10% of TSS efficiency variation.

**Figure 6.**
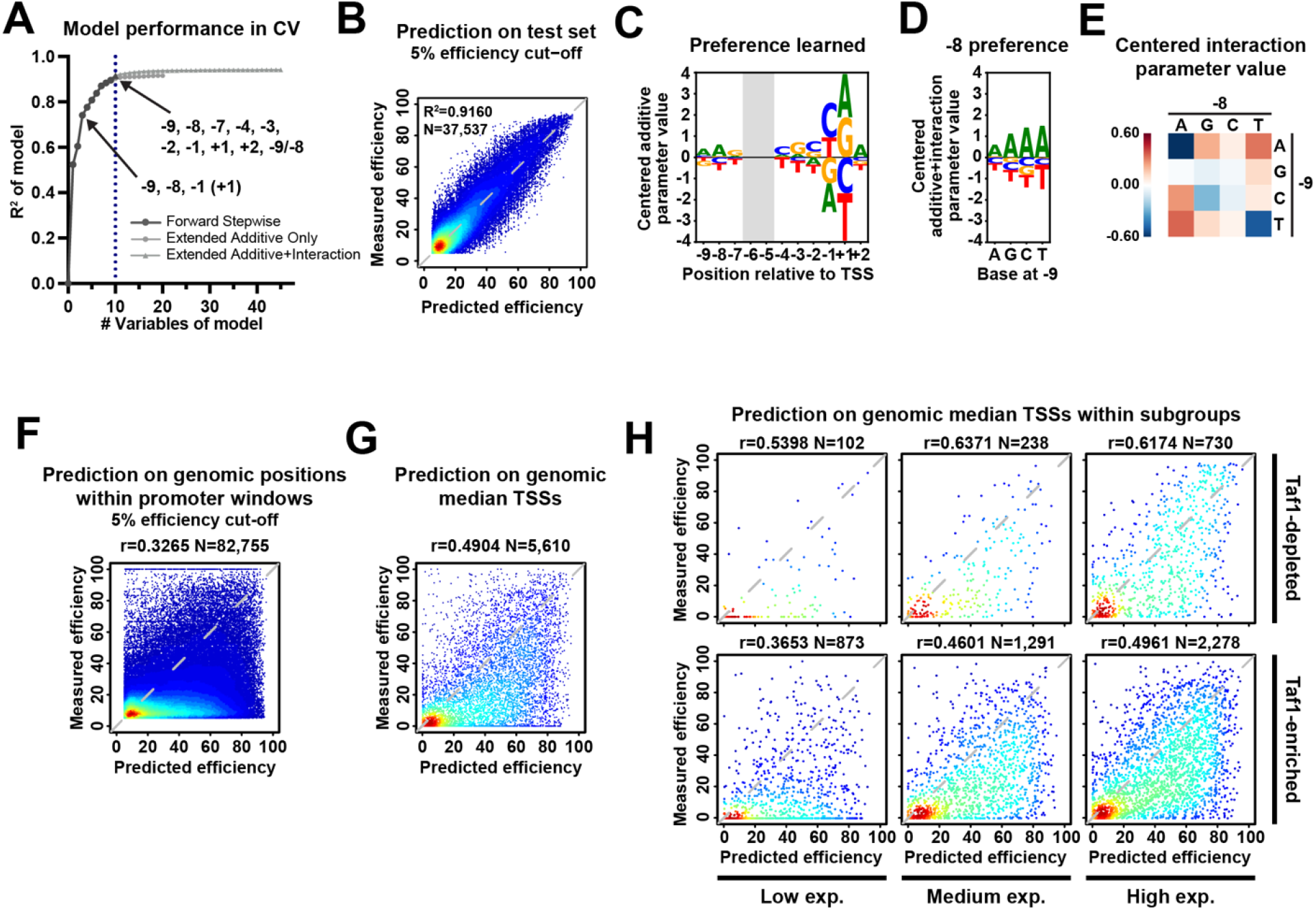
Logistic regression model of DNA sequence contribution to TSS efficiency. **(A)** Regression modeling identifies key DNA sequences and interactions contributing to TSS efficiency. Nine additive parameters plus one interaction were selected for the final model. Dots represents average R^2^ obtained in a 5-fold Cross-Validation (CV) strategy for logistic regression models using different numbers of features. The black line with SD error bars represents models with the best performance under a certain number of predictors. Note that model with as few as three additive parameters could explain 74.10% of TSS efficiency variation in our WT libraries. **(B** to **E)** Good performance of model including sequences at nine positions and the -9/-8 interaction indicates that TSS efficiency in our libraries is mainly regulated by the included features. (**B**) A scatterplot of comparison of measured and predicted efficiency of test sets, with a 5% efficiency cut-off. Model performance R^2^ on entire test and number (N) of data points shown in plot are shown. (**C**) A sequence logo of centered additive parameters. The coefficients for bases at a particular position was centered and visualized as a sequence logo. (**D**) A sequence logo showing learned preference at position -8 when different bases existing at position -9, with -9/-8 interaction included. The -9/-8 interaction parameters were added to corresponding additive coefficients for bases at position -8. The additive plus interaction parameters were then centered and visualized as sequence logos. (**E**) A heat map of centered parameters for -9/-8 interaction illustrating how bases at one position affect preference at another position. **(F** to **G)** Efficiency prediction for positions within known promoter windows in WT shows overall over-prediction. Scatterplots of comparison of measured and predicted TSS efficiencies of all positions (with a 5% efficiency cut-off) (**F**) or median TSSs (**G**) within 5979 known genomic promoter windows ^21^ with available measured efficiency. **(H)** Model shows better performance on Taf1-depleted promoters and promoters with medium to high expression. Scatterplots of comparison of measured and predicted TSS efficiency of median TSSs subgrouped by promoter classes and expression levels. Expression levels of genomic promoters are defined based on their total TSS-seq reads in the promoter window in the examined datasets: low, [0, 200); medium, [200, 1000); high, [1000, max). Pearson r and number (N) of compared variants are shown.

The final model containing the most predictive features explained 91.60% of the variance in TSS efficiency for WT test set (20% of total dataset) (**Figure 6B, Figure S5B**). Principal Component Analysis (PCA) on variables of models trained with individual replicates of WT and Pol II mutants showed large differences between WT and Pol II mutants but very small differences between replicates (**Figure S5C**), indicating that modeling captured features of different Pol II activity groups. We next asked whether the features learned by modeling using compiled datasets were consistent with our previous sequence preference analysis using selected representative datasets with the most randomized bases. We centered additive variable values and visualized results as a sequence logo (**Figure 6C**). First, as expected, positions -1 and +1 were the major predictors, however the influence of the -8 A did not appear as strong as in our previous preference analysis. We suspected this might be because the -9/-8 interaction contribution was not included. After adding the -9/-8 interaction term, we observed emergence of the position -8 as an influential predictor (**Figure 6D, E**), which also emphasizes the contribution of the -9/-8 interaction. Second, and importantly, modeling confirmed the +2A preference observed in previous motif enrichment analysis using only the most efficient -8 TSS variants (**Figure S2D**). The impact of position +2 is also evident when performing Principal Component Analysis (PCA) to variables of models trained with WT or Pol II mutant datasets (**Figure S5C**). The fact that sequences at position +2 are the top contributing variables in 2^nd^ principal component that distinguished G1097D and E1103G is in agreement with differentially altered +2 sequence enrichment by Pol II mutants (**Figure S3I**). These results suggest position +2 preference is altered by Pol II activity changes, and that this position might work directly with the Pol II active site.

### Sequence defines TSS efficiency within a wider promoter context during initiation by scanning

To evaluate the extent to which DNA sequence around a TSS contributes to TSS efficiency at genomic promoters, we compared the differences between observed and model-predicted efficiencies for all positions within genomic promoter windows or within specific subgroups of known promoters (**Figure 6F-H, Figure S5D**). As expected, we found most promoter positions showed low or no observed efficiency and were over-predicted by sequence alone (**Figure 6F, Figure S5D**). This is because TSSs need to be specified by a core promoter followed by scanning within a limited range. Therefore, features beyond TSS sequence, such as distance from the site of PIC assembly, determine initiation in the genome. We therefore extracted observed median TSSs as representatives of TSSs in contexts supporting efficient initiation to ask how our sequence-based predictor functioned on genomic TSSs (**Figure 6G**). We also separated median TSSs by promoter classes based on Taf1 enrichment ^68^, a proxy for the two main types of promoters in yeast, and by promoter expression levels (**Figure 6H**). We observed good prediction performance for a wide range of TSSs indicating that sequence determinants identified in our limited promoter context also contribute to TSS efficiency in genomic promoter contexts. We observed increased performance for more highly expressed promoters (Pearson r increased from 0.37-0.54 to 0.46-0.64) and for Taf1-depleted promoters (Pearson r increased from 0.37-0.50 to 0.54-0.64) (**Figure 6H**). Conceivably, highly expressed promoters may have evolved TSSs at optimal distances from core promoters, and therefore may be similarly sensitive to sequence effects. Alternatively, our libraries used *GAL1* UAS and *SNR37* core promoter elements. Since both *GAL1* and *SNR37* are highly-expressed, Taf1-depleted promoters, our libraries may share sequence sensitivities with TSSs from promoters with related architectures.

## Discussion

Here we developed and employed Pol II MASTER to systematically investigate ∼1 million TSS sequences in wildtype or Pol II mutant cells. This system allowed us to specifically and comprehensively study TSS efficiencies in initiation by promoter scanning by removing confounding effects from other architectural features, such as variability in core promoter-TSS distances, differences in promoter identities or chromatin configurations that may obfuscate analyses of genomic TSSs. We find that sequence variation at different positions around TSSs considerably tunes initiation efficiency in a predictable way and these contributions are important for initiation efficiency at genomic promoters.

Combining results from this study and others, we suggest how TSS selection during promoter scanning works through TSS sequence and Pol II activity (**Figure 7**). We find that two major sequence position groups contribute to TSS selection: bases around the TSS and bases around position -8. First, in promoter scanning, the TSS and adjacent bases interact with Pol II active site, the 1^st^ NTP or each other to facilitate stable binding of 1^st^ NTP and potentially the 2^nd^ NTP to stimulate RNA synthesis. Sensitivity of initiation efficiency to NTP levels for the first through fifth transcription positions in vivo supports this idea. This function of downstream positions would be in contrast to the concept of the initiator or downstream elements functioning as part of the TFIID or PIC binding site, as has been proposed for higher eukaryotes (see ^69, 70^ and references therein). As the universal initiating element, Y_-1_R_+1_ has been established to facilitate stable binding of the NTPs by RNA polymerases via base stacking between R_-1_ from template DNA and the 1^st^ purine NTP ^43, 44^. Positions upstream of the TSS, such as positions -4 to -2, might contribute to stabilize template DNA via base stacking or physical interaction with initiation factors ^44, 71^.

**Figure 7.**
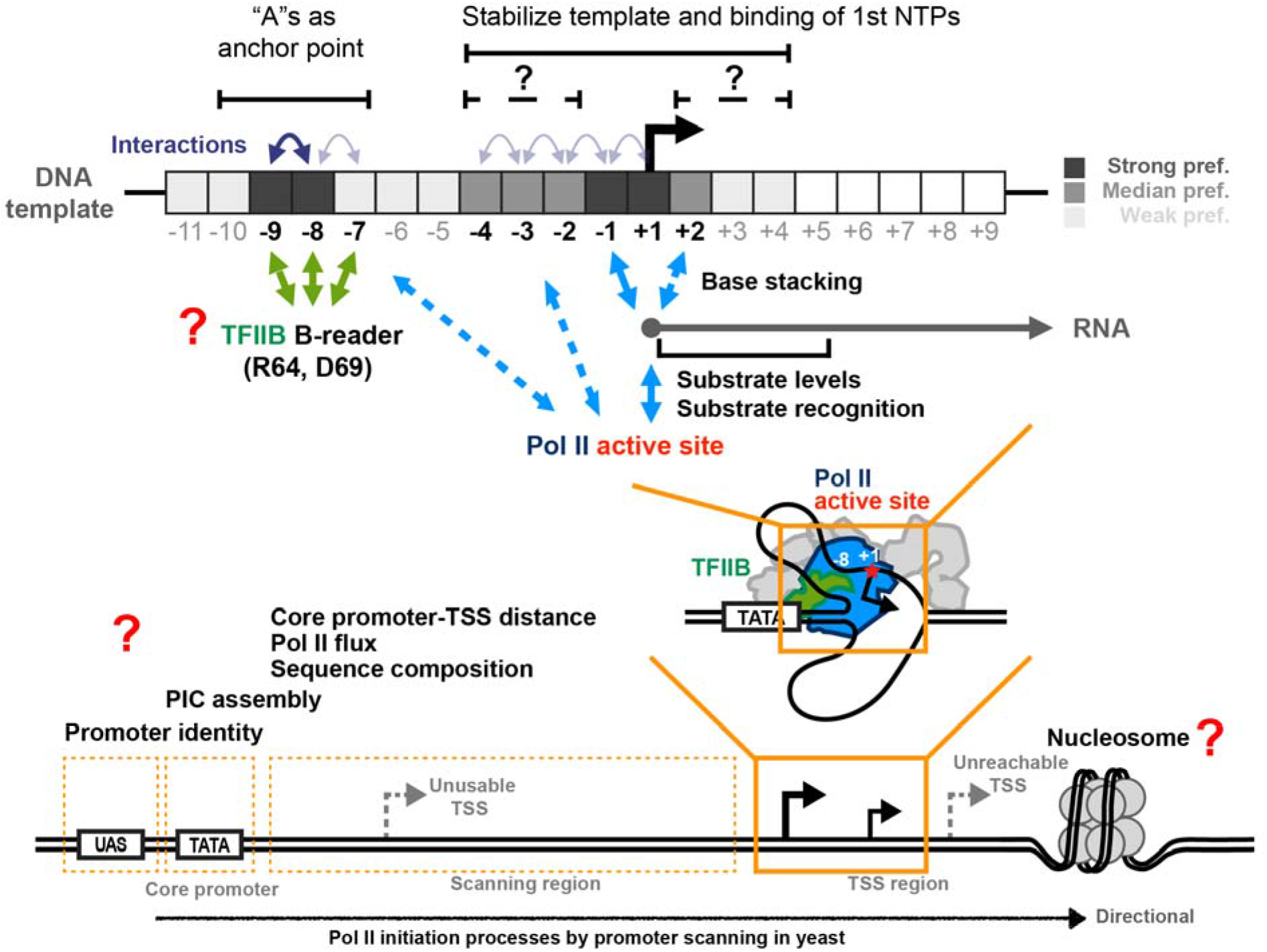
Model for TSS sequence preference regulated by multiple mechanisms. Top panels show determined contribution of sequence at positions around TSS and proposed mechanisms. Two major groups of positions around TSS contribute to TSS selection: bases around TSS (actual initiating site) and bases around position -8. The TSS and adjacent bases interact with Pol II active site, the 1^st^ NTP or each other to facilitate stable binding of 1^st^ NTP thus stimulate RNA synthesis. -8 and -9 Ts on the template strand with an additional interaction between -8 and -7 template strand positions may serve as an anchor point interacting with TFIIB B-reader domain allowing pausing of scanning and promotion of Pol II initiation at TSSs a fixed distance downstream, as proposed ^47^. These preferences are reflected as As if the analysis is on the coding strand. Positions and interactions that were identified by regression modeling as robust features are labelled in bold. Bottom panel shows other architectural features involved in Pol II transcription initiation likely additionally contributing to TSS selection and initiation efficiency that will be accessible to Pol II MASTER analysis.

Our observation that Pol II mutants showed selective effects on based on initiating NTP (ATP or GTP specified by TSS position +1) (**Figure 3**) supports that +1 NTP efficiency is directly sensitive to properties of the Pol II active site during initiation by promoter scanning. Meanwhile, the observation that Pol II initiation is sensitive to NTP pools (**Figure 4C, Figure S4A**) further suggests a mechanism for cellular state to regulate initiation via alteration of NTP levels for the first position and several subsequent transcript positions. For GOF Pol II mutants, we propose that differential preference for ATP versus GTP is a direct consequence of Pol II active site changes. Pol II has been observed to bypass cyclobutane pyrimidine dimers (CPD) in vitro through addition of an untemplated A across from CPD lesions and a GOF Pol II appears to promote this activity, suggestion a potential increase in selectivity for ATP in some situations ^72^. In contrast to effects observed for GOF mutants, we propose that tested LOF mutants may have selective effects on TSS efficiency for +1A versus +1G sites due to indirect effects on NTP levels in cells. Such defects might result from altered synthesis of nucleotide synthesis-related genes, a number of which are themselves sensitive to Pol II activity ^23, 67, 73–76^. We observed the LOF mutant F1086S indeed alters all NTP levels (**Figure 4A**). Specifically, the LOF mutant increased GTP level, which may be due to the constitutive expression of *IMD2* in Pol II LOF mutants ^23, 74^, and the overall increase in expression for *IMD* genes at the mRNA level ^23^.

While levels of NTP substrates are expected to impinge on initiation at some point, it is striking that modest reduction or increase in GTP, CTP, or UTP levels appear to shape initiation efficiency genome-wide. These changes suggest that environmental control of initiation may be used more broadly than anticipated and that distinct transcripts will be differentially affected upon perturbation of NTP levels. Direct sensing and regulation of promoter activity by NTP levels has been proposed for the *IMD2* promoter in yeast ^67, 77^. At this promoter, upstream +1G TSSs are used to generate a non-productive set of transcripts that terminate before transcription of the *IMD2* reading frame under GTP-replete conditions. Upon GTP limitation (or catalytic defects in Pol II), productive initiation is observed at a downstream +1A TSS. Our results here argue that nucleotide sequence content is sufficient to confer the observed regulation as we find that most +1G TSSs in the genome are reduced for initiation efficiency upon reduction in GTP levels. For select TSSs from *IMD2* that they tested, the Brow lab had previously found that Gs at the first or second transcript positions could confer MPA sensitivity to a TSS ^67^. Here, we find this effect genome-wide and that it extends to transcript +5 position for GTP while finding evidence for effects of UTP and CTP levels.

Consistent with NTP effects described above, positions +2 to +4 downstream of TSS could contribute by establishish initial RNA and NTP stability in the Pol II active site (**Figure S2D**). We observed Pol II mutant effects on those preferences (**Figure S3I, Figure S5C**), suggesting these positions might function via directly interacting with the Pol II active site, as observed in transcription structures in other species ^43, 44^. First, the T7-like single-subunit RNAP family shows a base-specific interaction between the +2 NTP and a residue in the middle of the O helix, which was suggested to enhance formation of the first phosphodiester bond ^43^. Second, a structure of *de novo* transcription initiation complex in bacterial RNA Polymerase showed multiple interactions between the 2^nd^ NTP and its β’ and β subunits, whose eukaryotic counterparts are Rpb1 and Rpb2 ^44^. Alternatively, the favored A/G-rich A(/G)_+1_A(/G)_+2_G_+3_G_+4_ motif might be related to efficiency of enzyme translocation. A study from the Landick lab showed that when A/G comprised the RNA 3’ end, the RNAP active site favored the post-translocated state ^78^. If A/G similarly affects translocation state within first four bases in Pol II initiation, synthesis of the first few bases might be promoted. Our results detecting sensitivity in initiation efficiency based on levels of NTPs specified at positions downstream from +1 (**Figure 4, S4**) suggest potential abortive initiation under conditions that slow phosphodiester bond formation early in transcript synthesis. Abortive initiation by yeast Pol II in vitro under NTP replete-conditions is very low in specific artificial initiation conditions ^79^. However, these conditions did not involve initiation by a scanning PIC with all GTFs or reduced NTP levels. Revisiting the biochemical properties of Pol II during bona fide initiation by promoter scanning could be valuable and our studies make specific predictions that can be tested.

The GTF TFIIB has been specifically implicated in the preference for bases near position -8, because the TFIIB B-reader domain has been observed to directly interact with a template T at -8 in a structure of a yeast Pol II-TFIIB complex ^80^. Here, it is attractive to envision TFIIB functioning as an anchor point allowing pausing or slowing of the scanning process to promote Pol II initiation at a fixed distance downstream. Several observations support this proposed function. First, we detected sequence interaction between positions -8 and -7 (**Figure 2G, Figure S2F**). This is in line with the direct contact of -7T and -8T on the template strand and TFIIB B-reader R64 and D69 observed in Pol II-TFIIB complex structure ^80^. Second, we observed a strong interaction between positions -9 and -8, where the presence of an A at either position suppressed the preference of A at the other position (**Figure 2F, Figure S2E**). This -9/-8 interaction was also evident when examining genomic median TSSs (**Figure 5C**). Taken together, we speculate that Ts around position -8 or -9 on the template strand and TFIIB may pause the scanning process to facilitate the usage of TSSs positioned 8 to 9 bases downstream. Moreover, we have shown that Pol II catalytic mutants alter TSS efficiencies across all TSS sequences, without showing alteration in preference for -8A (**Figure 3, Figure S3F**), suggesting that Pol II catalytic activity is not responsible for the -8 preference.

Whether or how DNA sequence surrounding TSSs is involved in other promoter properties is another question. “A” bases at positions -7 to -5 were observed to be neutral in our promoter libraries (**Figure 2E**), in contrast to the apparent A-enrichment observed for highly expressed and focused genomic TSSs (**Figure 5A, B**) ^1, 16, 21, 34, 45^. We speculate that observed A-richness around TSS functions through other evolved promoter properties. <u>First</u>, the observed A-richness between positions -10 to -3, together with T-richness at further upstream core promoter regions ^45, 46^, provides an easily meltable region for DNA unwinding, perhaps facilitating transcription initiation in specific contexts. <u>Second</u>, higher A/T content may function to lower nucleosome occupancy ^81^, because appropriately periodic G/C dinucleotides promote nucleosome occupancy ^82–85^. However, the base composition switch in highly expressed promoters from T-to A-preponderance ^21, 45, 86^ indicates A-richness may have other roles depending on the characteristics of the sequence itself. <u>Third</u>, A-richness may be left over from the evolution of promoter scanning. A recent study of transcription initiation mechanism investigated 12 yeast species and proposed that during evolution a A-rich region upstream of TSS appeared first, then the specific -8A preference occurred ^47^. Therefore, the observed A preference at upstream positions of highly used and focused TSSs may be leftovers of A-enrichment in those promoters during scanning evolution in addition to promoter roles beyond TSS selection *per se*.

Our studies highlight the strength of approaches that minimize contextual factors by isolating specific promoter attributes for study in high-throughput. Here we have employed Pol II MASTER to the DNA sequence determinants of initiation efficiency during Pol II scanning. It will be valuable to apply this systematic analysis to other promoter architectural factors determining Pol II initiation, such as UAS identity, core promoter-TSS distance and sequence composition within the scanning region. In addition, we have found sequences downstream of TSSs contribute to TSS efficiency. Therefore, expanding the randomized region is needed to refine our understanding of sequence preference and potential longer range sequence interactions. Furthermore, applying Pol II MASTER across initiation mutants and promoter variants will reveal factor-sequence relationships and may allow initiation potential to be determined from DNA sequence and genome location alone.

## Methods

### Yeast strains, plasmids, oligonucleotides and media

Yeast strains, plasmids and oligonucleotide sequences are described in **Table S1**. All oligonucleotides were obtained from IDT. Yeast strains used in this study were constructed as previously ^21, 23, 73, 87^. Briefly, plasmids containing *rpb1* mutants (G1097D, E1103G, F1086S, and H1085Q) were introduced by transformation into yeast strain CKY749 containing a chromosomal deletion of *RPO21/RPB1* but with a wild type *RPB1 URA3* plasmid, which was subsequently lost by plasmid shuffling. All plasmids and yeast strains are available upon request. Yeast media are following standard protocols ^88^. YPD solid medium is made of yeast extract (1% w/v; BD), peptone (2% w/v; BD, 211677), bacto-agar (2% w/v; BD, 214010) and dextrose (2% w/v; VWR, VWRBK876) supplemented with adenine (0.15 mM; Sigma-Aldrich, A9126) and L-tryptophan (0.4 mM; Sigma-Aldrich T0254). Minimal media plates are synthetic complete (“SC”) with amino-acids dropped out as appropriate as described in ^88^ with minor alterations as described in ^23^: per standard batch formulation, adenine hemisulfate (Sigma-Aldrich, A9126) was 2 g, L-Leucine (Sigma-Aldrich, L8000) was 4 g, myo-inositol was 0.1 g, para-aminobenzoic acid (PABA) was 0.2 g.

### Construction and transformation of plasmid libraries

A 9 nt randomized TSS region and 20 nt randomized barcodes, with 4 fixed bases inserted between every 4 nt (NNNNANNNNCNNNNTNNNNGNNNN), were separately synthesized by IDT as oligo pools with specific randomized positions using “hand mixing” for N positions to ensure even randomization and avoid bias during machine mixing of precursors during oligo synthesis. Together with other components including the *GAL1* UAS, *SNR37* core promoter, *SNR37* TSS region (“flux detector”), *GFP* ORF, and the *CYC1* terminator, template libraries were constructed by PCR sewing and cloned into pRSII413 (a gift from Steven Haase, Addgene plasmid #35450; http://n2t.net/addgene:35450; RRID:Addgene_35450) ^89^ by ligation (**Figure S1A**). Ligation products were transformed into *Escherichia coli* TOP10F’ cells (Invitrogen) and grown on LB plates supplemented with carbenicillin (100 µg/ml) at high density. 200,000-500,000 colonies were collected from each library to maximize variant representation. Plasmid libraries were isolated from cells pellets using ZymoPURE II Plasmid Maxiprep Kit (Zymo Research, D4203) per manufacturer’s instructions. Plasmid library pools were transformed into yeast strains with wildtype and mutated Pol II using chemical transformation and electroporation, respectively. For Pol II WT libraries, 500 ng plasmid pool per reaction was transformed following yeast high efficiency transformation protocol described in ^90^. For Pol II mutant libraries, 2 µg plasmid pool per reaction was electroporated into Pol II mutant strains following yeast electroporation transformation protocol described in ^91^, with 50 µg single-stranded carrier DNA added. Transformants were grown on selective SC-His plates with 2% glucose as carbon source at high density. Three biological replicates were performed for each library and on average over two million colonies were collected for each replicate. Transformants scraped from densely plated transformation plates were inoculated into fresh SC-His medium with 2% raffinose (Amresco, J392) at 0.25 x 10^7^ cells/ml and grown until 0.5-0.8 x 10^7^ cells/ml, as determined by cell counting. Subsequently, galactose (Amresco, 0637) was added for three hours (4% final concentration) to induce library expression. For MPA treated WT libraries, MPA (in 100% ethanol; Sigma-Aldrich, M3536) was added for 20 min (20 µg/ml final concentration) prior to the three-hour galactose induction, with same volume of 100% ethanol as solvent control. 50 ml and 5 ml culture aliquots, for RNA and DNA extraction respectively, were harvested and then cell pellets were stored at -80 °C for further processing as described below.

### Generation of DNA amplicons for DNA-seq

Plasmid DNA from yeast cell pellets was isolated using YeaStar Genomic DNA Kit (Zymo Research, D2002) per manufacturer’s instructions. Amplicon pools containing the TSS and barcode regions were generated using plasmid DNA from *E.coli* or yeast by Micellula DNA Emulsion & Purification (ePCR) Kit (EURx/CHIMERx, 3600) per manufacturer’s instructions. To minimize amplification bias, each sample was amplified in a 15-cycle ePCR reaction, purified and subject to an additional 10-cycle scale-up ePCR reaction. To create the necessary sequence template diversity for Illumina sequencing, 18-25 bp and 1-7 bp “stuffer” sequences were added to 5’- and 3’-ends, respectively, during amplicon preparation. Amplicon pools were subject to Illumina NovaSeq 6000 (150 PE) sequencing, and on average 20 M paired-end reads were obtained from each replicate of a sample, with high reproducibility and minimal perturbation of the variant distribution with each library (**Table S2**).

### Sample preparation for TSS-seq

Total RNA was extracted by a phenol-chloroform method ^92^, followed by RNA purification (RNeasy Mini kit, QIAGEN, 74104) with on-column DNase digestion (RNase-Free DNase Set, QIAGEN, 79254) to remove DNA. TSS-seq was done using procedures described in ^93^. To prepare RNAs for the cDNA library construction, samples were sequentially treated with Terminator 5‘-Phosphate-Dependent Exonuclease (Lucigen), Quick CIP (calf-intestine alkaline phosphatase, NEB) and Cap-Clip™ Acid Pyrophosphatase (CellScript) to remove 5’ monophosphate RNA and convert 5’ triphosphate or capped RNAs to 5’ monophosphate RNAs. Next, RNA prepared with enzymatic treatments was ligated to the 5’-adapter (s1206-N15, 5’-GUUCAGAGUUCUACAGUCCGACGAUCNNNNNNNNNNNNNNN-3’) that contains Illumina adapter sequence and a 15 nt randomized 3’-end to reduce ligation bias and serve as a Unique Molecular Identifier (UMI). Next, cDNA was constructed by reverse transcription using RT primer CKO2191-s128A (5’-CCTTGGCACCCGAGAATTCCAAGTGAATAATTCTTCACCTTTA-3’) followed by emulsion PCR amplification for 20-22 cycles using Illumina PCR primers (RP1 and RPI3-30). Final DNA was gel size selected for 180-250 bp lengths and sequenced by Illumina NextSeq 500 (150 SE) or NovaSeq 6000 (200 SE) (**Table S2**) using custom primer s1115 (5’-CTACACGTTCAGAGTTCTACAGTCCGACGATC-3’) to avoid potentially confounding effects of misannealing of the default pooled Illumina sequencing primers to the two randomized sequence regions.

### Primer extension assay

Primer extension assays were performed on the same batch of total RNA extracted for TSS-seq as described in ^94^ with modifications described in ^23^. For each reaction, 30 µg total RNA was used. An RNA sample without library transformed was used as “no GFP” control. A sample containing same amount of nuclease-free water was used as “no RNA” control. A primer (CKO2191) complementary to the 6^th^ to 27^th^ bases of *GFP* ORF, which is the same annealing region for reverse transcription of TSS-seq sample preparation, was labeled with ^32^P γ-ATP (PerkinElmer, BLU502Z250UC) and T4 polynucleotide kinase (Thermo Scientific, EK0031). M-MuLV Reverse Transcriptase (NEB, M0253L), RNase inhibitor (NEB, M0307L), dNTPs (GE) and DTT were added to mix of RNA and labelled primer for reverse transcription reaction. Before loading to sequencing gel, RNase A (Thermo Scientific, EN0531) was added to remove RNA. The products were analyzed by 8% acrylamide/bis-acrylamide (19:1 ratio, Bio-Rad, 1610145) gel containing 1x TBE and 7M Urea. Primer extension gels were visualized by a Molecular Imager PharosFX™ Plus System (Bio-Rad) and quantified by Image Lab (5.2).

### Determination of NTP levels

For each genotype or treatment, six biological replicates were performed. Cells from saturated overnight cultures were used to inoculate fresh SC medium containing 2% raffinose at 0.25 x 10^7^ cells/ml and grown to a density of 0.5-0.8 x 10^7^ cells/ml, as determined by cell counting. For each WT replicate, two 15 ml cultures were aliquoted and treated with 20 ug/ml MPA or 100% ethanol for 20 min, respectively. Subsequently, three-hour 4% galactose treatment was performed for all samples. About 1 x 10^7^ cells were harvested, and then cell pellets were snap frozen using liquid nitrogen and immediately stored at -80 °C for further processing as described below.

Metabolic quenching and nucleotide phosphate metabolite pool extraction were performed by adding 1 ml ice-cold 2:2:1 acetonitrile:methanol:water with 0.1% formic acid with 1x MS-Stop phosphatase inhibitor (Sigma-Aldrich). Samples were spiked with 10 µl 100 mM mix of adenosine-^15^N_5_-5′-triphosphate, guanosine-^15^N_5_-5′-triphosphate, thymidine-^13^C_10_,^15^N_2_-5′-triphosphate, cytosine-^2^H_14_-5′-triphosphate, and uridine-^15^N_2_-5′-triphosphate (Sigma-Aldrich). Samples were homogenized via sonification at room temperature, and the supernatant was then cleared of protein by centrifugation at 16,000 xg. The protein pellet was resuspended in RIPA buffer for protein normalization. The cleared supernatant was dried down under nitrogen gas and resuspended in 100 µl 7.5 mM ammonium acetate / 0.05% ammonium hydroxide. 2µl of cleared supernatant was subjected to online LC-MS analysis. Purified adenosine-5′-triphosphate, guanosine-5′-triphosphate, thymidine-5′-triphosphate, cytosine-5′-triphosphate, and uridine-5′-triphosphate (Sigma-Aldrich) were serially diluted from 50 pmol/µl to 0.39 pmol/µl to generate calibration curves.

Analyses were performed by untargeted LC-HRMS. Briefly, samples were injected via a Thermo Vanquish UHPLC and separated over a reversed phase Phenomenex Kinetix-Polar C18 column (2.1 x 100 mm, 1.7 μm particle size) maintained at 55°C. For the 22.5-minute LC gradient, the mobile phase consisted of the following: solvent A (water/7.5 mM ammonium acetate/0.05% ammonium hydroxide) and solvent B (acetonitrile/0.05% ammonium hydroxide). The gradient was the following: 0-.1 min 2% B, increase to 70% B over 12 minutes, increase to 98% B over 0.1 min, hold at 98% B for 5 minutes, re-equilibrate at 2% B for five minutes. The Thermo IDX tribrid mass spectrometer was operated in positive ion mode, scanning in ddMS^2^ mode (2 μscans) from 200 to 800 m/z at 120,000 resolution with an AGC target of 2e5 for full scan, 2e4 for ms^2^ scans using HCD fragmentation at stepped 15,35,50 collision energies. Source ionization setting was 3.0 kV spray voltage for positive mode. Source gas parameters were 35 sheath gas, 12 auxiliary gas at 320°C, and 8 sweep gas. Calibration was performed prior to analysis using the Pierce^TM^ FlexMix Ion Calibration Solutions (Thermo Fisher Scientific). Integrated peak areas were then extracted manually using Quan Browser (Thermo Fisher Xcalibur ver. 2.7). Calibration curves using purified standards were then used to convert peak area ratios to concentration.

### Computational analyses

Data and statistical analyses were performed in Python (3.8.5) and R (4.0.0) environments. Additional packages usages are reported throughout the methods description. Source code is provided at https://github.com/Kaplan-Lab-Pitt/PolII_MASTER-TSS_sequence. Raw sequencing data have been deposited on the NCBI SRA (Sequence Read Archive) under the BioProject accession number PRJNA766624. Processed data have been deposited on the GEO (Gene Expression Omnibus) under the accession number GSE185290. Visualizations were compiled in Adobe Illustrator 2021.

#### DNA-seq analysis

High-throughput sequencing of template DNA amplicon was used to assign each 9 nt randomized TSS sequence to a corresponding 24 nt barcode. First, paired-end reads were merged using PEAR (0.9.11) ^95^. Next, we considered only those reads that contained a perfect match to three sequence regions common to all variants: 27 nt sequence upstream of the TSS region, 24 nt sequence between TSS region and barcode, and 27 nt sequence downstream of barcode (5’-TTCAAATTTTTCTTTTGATTTTTTTTC<u>NNNNNNNNN</u>ACATTTTCAAAAGGCTAACATC AG<u>NNNNANNNNCNNNNTNNNNGNNNN</u>ATGTCTAAAGGTGAAGAATTATTCACT-3’, randomized TSS and barcode regions are underlined). From these reads, 9 nt TSS region and 24 nt barcode were extracted, followed by individually error correction using UMI-tools (1.0.0) ^96^. Next, for barcodes linking to multiple TSS variants, only barcodes for which >= 90% of the sequencing reads containing a specified barcode also contained a shared, exact 9 nt TSS region were kept. To generate a master pool of TSS-barcode linkages for all TSS-seq samples, for each library (“AYR”, “BYR”, “ARY”), TSS-barcode linkages that existed in at least two out of four samples (one *E.coli* sample plus three WT yeast replicates) and in which >= 5 reads existed were kept and pooled. Two types of processed data are available in GEO database, with accession numbers listed in (**Table S2**): tables containing TSS-barcode linkages and corresponding DNA-seq read counts for each sample, tables of the master pool containing kept TSS-barcode linkages and corresponding DNA-seq read count in all related samples.

#### TSS-seq analysis for libraries

High-throughput sequencing of RNA samples was used to link RNA products to barcodes, therefore assign TSS usage to corresponding DNA templates. For TSS identification and subsequent preference analysis, we considered only those reads that contained a perfect match to a 27 nt sequence region downstream of the barcode, as well as expected length of 5’-end: 5’-[15 nt 5’-UMI]-[>1 nt upstream of barcode region, designated as “RNA 5’-end”]-[24 nt barcode]-[the first 27 nt of GFP ORF, ATGTCTAAAGGTGAAGAATTATTCACT]-3’. Those reads carrying one or more mismatches from expected sequences were used for analysis of relative expression of promoters. Next, 15 nt 5’-UMIs, “RNA 5’-end” with varying length, and 24 nt barcode were extracted and individually corrected by UMI-tools (1.0.0). Deduplication was performed based on 5’-UMIs, meaning reads contained a shared UMI-“RNA 5’-end”-barcode linkage were counted as one deduplicated read for further analysis. Next, the identity of the 24 nt barcode was used to determine the template sequences of randomized TSS region. Only reads with “RNA 5’-end” sequence perfectly matched to corresponding template sequence were used for analysis of TSS efficiency, but all deduplicated reads, including reads with mismatch(es), were used for analysis of relative expression. Next, a TSS-seq count table containing TSS usage distribution of each TSS promoter variant was generated. In the count table, each row represents one TSS promoter variant, and each column represents one position between positions -68 to +25 relative to “designed” +1 TSS. The number in each cell represents TSS-seq reads generated from a particular position, with perfectly match to DNA template. After investigating reproducibility (**Figure S3B-C, S4A-C**), count tables generated from three biological replicates were merged into one by aggregating read counts at each position. Promoter variants with >= 5 TSS-seq reads in each replicate and whose Coefficient of Variation (CV) of TSS-seq reads is <= 0.5 were kept. Two types of processed data are available in GEO database, with accession numbers listed in **Table S2**: tables containing “RNA 5’-end”-Barcode linkages and corresponding deduplicated TSS-seq read counts for individual sample, TSS-seq count tables for individual samples and after aggregating replicates. As an example, a TSS-seq count table including positions -10 to +25 relative to “designed” +1 TSS after aggregating replicates of WT libraries is shown as **Table S4**.

#### TSS efficiency calculation

TSS efficiency for each position was calculated by dividing reads count at a particular position by the reads at or downstream of this TSS. TSS positions with >= 20% efficiency but with <= 5 reads left for this and following positions were filtered out, as well as their downstream positions.

#### Relative expression analysis for libraries

The relative expression from each library promoter variant was defined as the ratio of normalized TSS-seq reads generated from a particular promoter variant to normalized DNA-seq reads containing the variant. The normalized DNA-seq or RNA-seq reads were calculated by dividing reads per promoter variant by the total number of reads per sample, followed by averaging three biological replicates. Promoter variants with >= 10 DNA-seq reads in each replicate and whose Coefficient of Variation (CV) of normalized DNA-seq reads is <= 0.5 were kept.

#### Sequence preference analysis

For sequence preference at each position, all TSS variants were subgrouped based on the bases at a particular position. TSS efficiencies of TSS variants were visualized as scatter plots using GraphPad Prism 9. Kruskal-Wallis with Dunn’s test was performed to test sequence preference in GraphPad Prism 9. Next, TSS efficiency medians of each subgroup were calculated and centered to calculate “relative efficiency” at each position. The relative efficiencies were visualized as sequence logos using Logomaker (0.8) ^97^. In motif enrichment analysis, surrounding sequences relative to examined TSSs were extracted and visualized as sequence logos using WebLogo 3 ^98^. Heatmaps, scatter plots and density plots for comparing Pol II WT and mutants were generated by Morpheus (https://software.broadinstitute.org/morpheus) or ggplot2 (3.3.3) R package.

#### Interaction analysis

The interaction between positions is defined as different bases existing at one position resulting in different sequence preferences at another position. For any two positions, all TSS variants were subgrouped based on bases at both positions. Median values of TSS efficiency distribution of each subgroup were calculated and centered twice to calculate “centered relative efficiency”. The centered relative efficiencies were visualized as heatmaps using Seaborn (0.11.0). Interactions related to positions between -11 to -9 were calculated using datasets of designed +4 TSS deriving from “AYR”, “BYR” and “ARY” libraries. Other interactions were calculated using datasets of designed +1 TSS deriving from “AYR” and “BYR” libraries.

#### TSS-seq analysis for genomic TSSs

Genomic TSS-seq datasets from our lab’s previous study were used for comparison of model to in vivo TSS usage ^27^. Quality control, read trimming and mapping were performed as described in ^21, 27^ to generate a TSS count table that contains TSS-seq reads at each individual position within known promoter windows (“median” TSS, 250 nt upstream and 150 nt downstream from median TSS position). TSS efficiency calculation and subsequent sequence preference analyses were performed as that for Pol II MASTER libraries. EtOH and MPA treatment for genomic TSSs are described in Results and these libraries were analyzed as TSS-seq was above. TSSs within a promoter were analyzed by their positions within the cumulative distribution of usage from upstream to downstream, with 50% of the distribution representing the median TSS position. To limit potential effects of marginal TSSs, analyses of MPA treatment were limited to TSSs showing ≥2% efficiency in the EtOH condition within the 25%-75% of the TSS distribution.

#### Prediction of TSS efficiency

To prepare datasets for modeling, positions of designed -8 to +2 and +4 TSSs of each promoter variant that have valid TSS efficiency were compiled as sequence variants. For each TSS variant, sequences at -11 to +9 positions relative to TSS, together with corresponding TSS efficiency, were extracted. 80% of dataset were randomly partitioned as training set and the rest 20% as testing set. To select robust features, a forward stepwise strategy with a 5-fold Cross-Validation (CV) was employed in two major stages, for additive terms and for interactions. Starting with no variable in the model, logistic regression models with one additional variable (the sequence at a particular position) were trained to predict TSS efficiency on training set by train() of caret (6.0.86) R package ^99^, with a 5-fold CV. The R^2^, representing the proportion of variance explained, was calculated to indicate the performance of each model. The variable that provides the highest increased R^2^ for model was added into the model for next round of variable selection. This process was repeated until the increased R^2^ is less than 0.01. After identifying the most influential additive variables, same process was repeated for investigating robust interactions between selected additive variables. Next, a final model with selected robust features. including additive variables and interactions, was constructed on entire training set using glm() and investigated on testing set. Comparison between predicted and measured efficiencies was visualized as scatter plots using LSD (4.1.0) R package. Model parameters were extracted and used to further calculation. Visualizations were done in Logomaker (0.8) and Seaborn (0.11.0) in Python. Principal Component Analysis (PCA) was performed using prcomp() in R.

## Supplementary Tables

**Supplementary Table 1.** Oligonucleotides, yeast strains, and plasmids.

**Supplementary Table 2.** Statistics and data accession information for DNA-seq and TSS-seq.

**Supplementary Table 3.** Summary of libraries.

**Supplementary Table 4.** TSS-seq count table of WT libraries.

## Supporting information

Supplemental Table 1 Strains Plasmids Oligos

Supplemental Table 2 Sequencing Information

Supplemental Table 3 Promoter Libraries Summary

Supplemental Table 4 TSSseq count table

## Acknowledgements

The authors thank Kaplan lab members for helpful comments on the manuscript. We are deeply grateful to Chenxi Qiu for discussions and comments on this project. We acknowledge Justin Kinney (Cold Spring Harbor Laboratory) and Shuoran Li (Statistical Consulting Center at University of Pittsburgh) for discussions on modeling. We thank Charles D. Johnson, Richard Metz (Texas A&M AgriLife Genomics and Bioinformatics Service), Andrew Hillhouse (Texas A&M Institute for Genome Sciences & Society), William A MacDonald, Rania Elbakri (the University of Pittsburgh Health Sciences Sequencing Core at UPMC Children’s Hospital of Pittsburgh), Yinghong Pan (the UPMC Genome Center), Dibyendu Kumar (the Waksman Genomics Core Facility at Rutgers University), and Liz Freeman (Illumina) for discussions and advice regarding deep sequencing strategies. We thank Steven J. Mullett and Stacy Gelhaus Wendell (Metabolomics and Lipidomics Core, NIHS10OD023402) for performing NTP measurements.

## Authors’ contributions

Y.Z. designed the project, performed experiments, analyzed data, made figures, drafted and revised the manuscript. I.V. generated libraries for TSS-seq. S-H.Z. analyzed data and discussed analysis. B.E.N. provided funding and methodology of TSS-seq, and revised the manuscript. C.D.K. conceived and designed the project, guided analyses and interpretation of data, provided funding, revised the manuscript.

## Data availability

Raw sequencing data generated in this study are available in the NCBI BioProject, under the accession number PRJNA766624. Processed data are available in GEO, under the accession number GSE185290.

## Code availability

Code for analyses in this study is provided at https://github.com/Kaplan-Lab-Pitt/PolII_MASTER-TSS_sequence.

## Figures and Figure Legends

**Figure S1.**
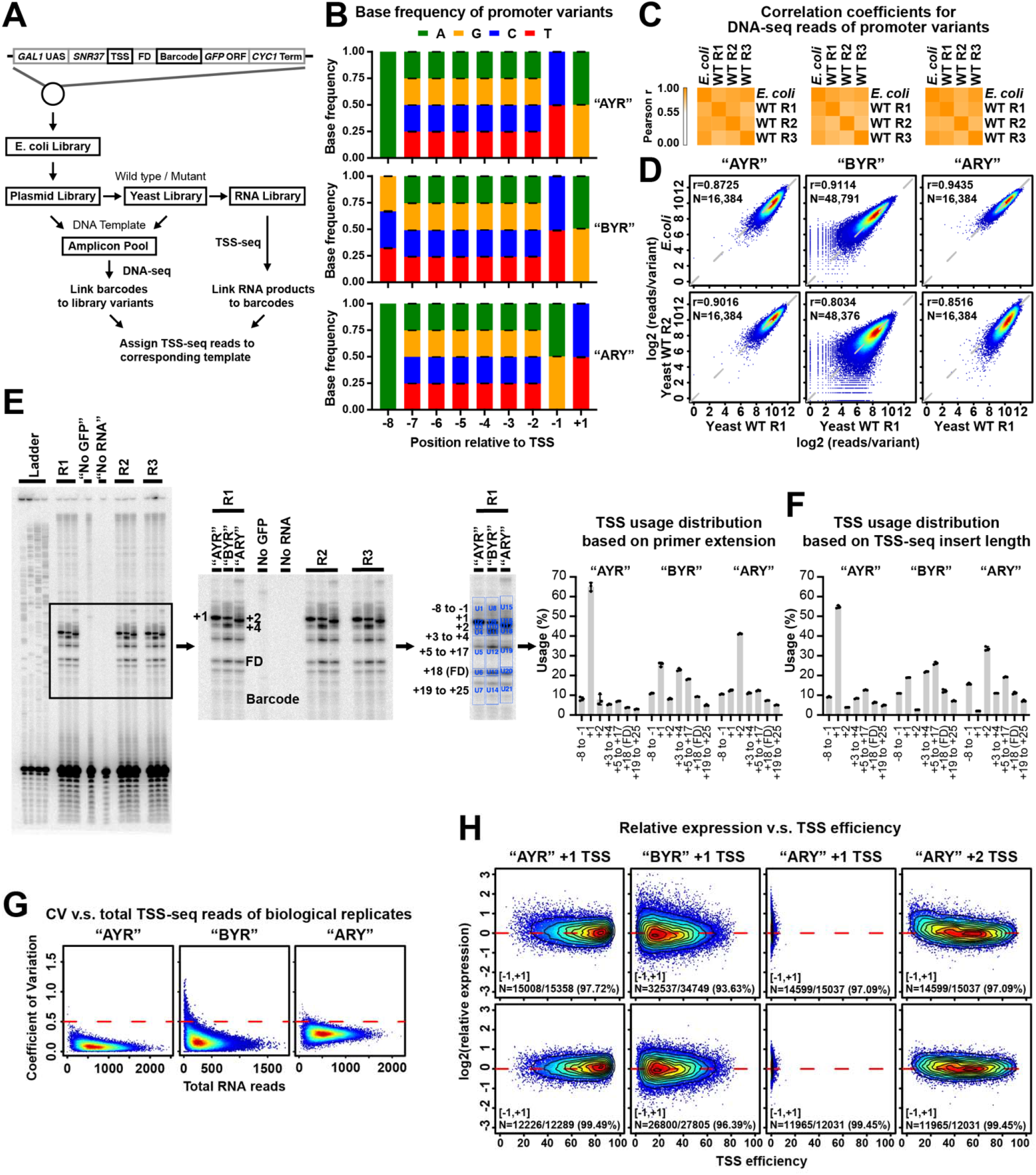
High level of reproducibility and coverage depth of library variants. **(A)** Schematic representation of experimental approach. Promoter libraries with almost all possible sequences within a 9 nt randomized region in a promoter context designed for specific functionalities were constructed on plasmids. Libraries were of three types, designated “AYR”, “BYR”, and “ARY” based on compositions of their randomized regions. Plasmids were amplified in *E. coli* and transformed into yeast strains with wild type or mutated Pol II. DNA and RNA were extracted from yeast pellets and prepared for DNA-seq and TSS-seq. **(B)** Base frequencies at positions within the randomized region of promoter variants demonstrate unbiased synthesis of randomized regions. Bars are mean +/-standard deviation of the mean for promoter variants in WT and four Pol II mutants. **(C)** Heatmap illustrating hierarchical clustering of Pearson correlation coefficients of reads per promoter variant for libraries amplified in *E. coli* and three biological replicates of libraries transformed into yeast. **(D)** Example correlation plots of DNA reads count of promoter variants for *E. coli* and yeast WT biological replicates. Pearson r and number of compared variants are shown. **(E)** Bulk primer extension for RNA produced from promoter variant libraries and quantification for biological replicates transformed into WT yeast. “No GFP” control used an RNA sample without a library transformed. “No RNA” control used a sample of nuclease-free water. Dots represent three biological replicates. Bars are mean +/-standard deviation of the mean. **(F)** TSS usage distribution based on insert length of TSS-seq reads generated from transformed libraries. Dots represent three biological replicates. Bars are mean +/-standard deviation of mean. Distributions are similar to the distributions in **(E)**. Note that primer extension will blur usage into adjacent upstream position due to some level of non-templated addition of C to RNA 5’ ends. **(G)** Heat scatter plots of Coefficient of Variation (CV, y axis) versus total RNA sequencing reads per promoter variant in each of three Pol II MASTER libraries. A cutoff of CV=0.5 was used to filter out variants with higher variance. **(H)** Heat scatter plots of relative expression versus TSS efficiency of major TSSs per promoter variant, with contour lines indicating deciles in the data. Number of promoter variants with [-1, +1] relative expression values (log_2_) and corresponding percentage of total promoter variants are shown.

**Figure S2.**
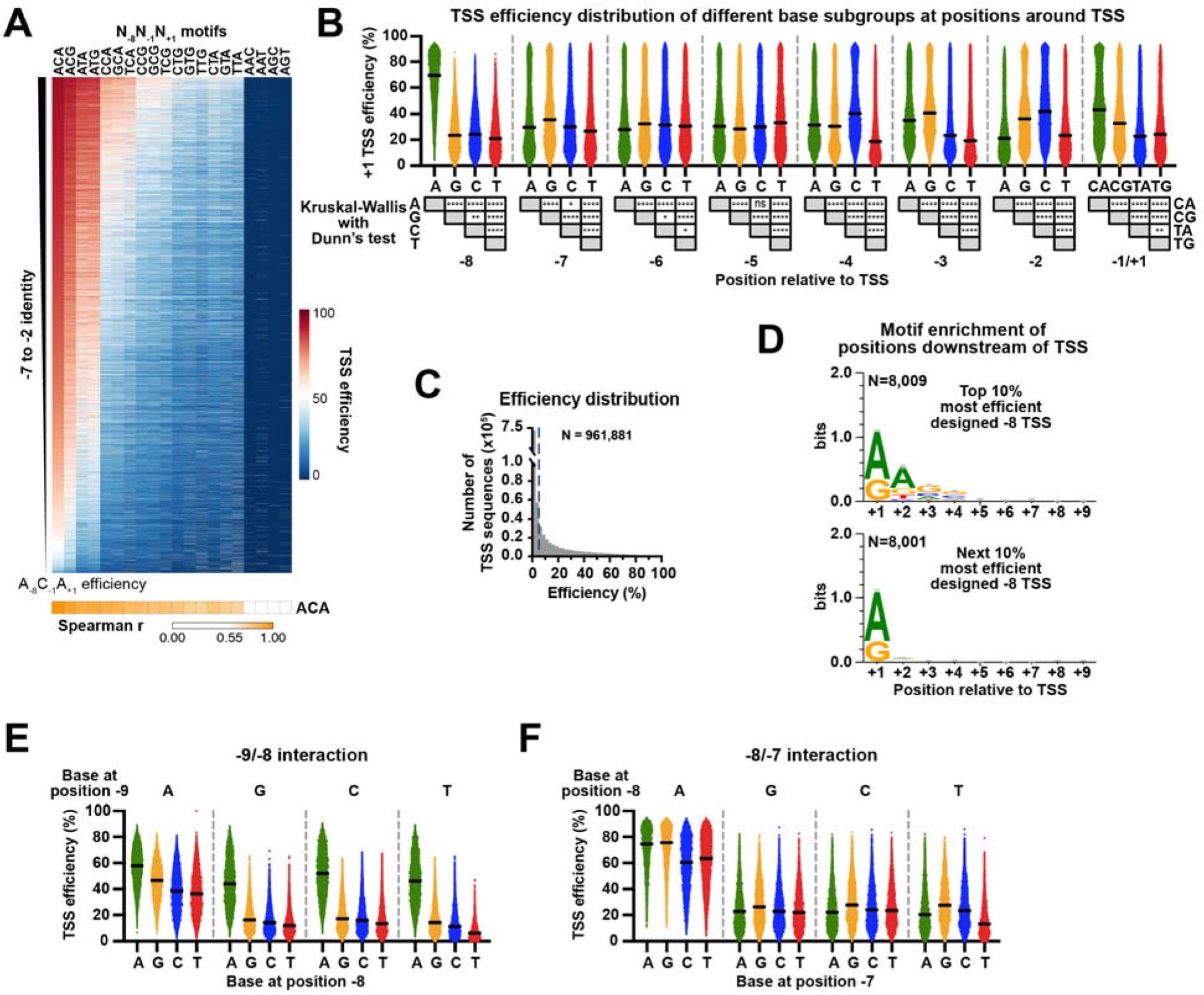
Surrounding sequence of TSSs modulates initiation efficiency. **(A)** +1 TSS efficiency of all -7 to -2 sequences within each N_-8_N_-1_N_+1_ motif in WT, rank ordered by efficiency of their A_-8_C_-1_A_+1_ version, is shown as a heat map. The x-axis is ordered based on median efficiency values for each N_-8_N_-1_N_+1_ motif group, as shown in **B**. Statistical analyses by Spearman’s rank correlation test between A_-8_C_-1_A_+1_ group and all groups are shown beneath the heat map. **(B)** Efficiency distributions of designed +1 TSSs grouped by base identities between -8 and +1 positions. Statistical analyses by Kruskal-Wallis with Dunn’s multiple comparisons test for base preference at individual positions relative to +1 TSS are shown beneath data plots. Lines represent median values of subgroups. ****, P ≤ 0.0001; ***, P ≤ 0.001; **, P ≤ 0.01; *, P ≤ 0.05. **(C)** Histogram showing the distribution of measured efficiencies for all designed -8 to +4 TSSs of all promoter variants from “AYR”, “BYR” and “ARY” libraries in WT. Dashed lines mark the 5% efficiency cutoff used to designate a TSS as active. Total number of TSS sequences is shown. **(D)** A_+2_G_+3_G_+4_ motif enrichment is apparent for the top 10% most efficient designed -8 TSS. A(/G)_+2_G(/C)_+3_G(/C)_+4_ motif enrichment was observed for the top 10% most efficient -8 TSSs but not for the next 10% most efficient TSSs. A(/G)_+1_ enrichment observed for top 20% most efficient TSSs is consistent with the +1R preference of TSS. Numbers (N) of variants assessed are shown. Sequence logos were generated using WebLogo 3. Bars represent an approximate Bayesian 95% confidence interval. **(E)** An A at position -9 results in different sequence preferences at position -8. The dataset of designed +4 TSSs deriving from “AYR”, “BYR” and “ARY” libraries was used to detect the -9/-8 interaction. All variants were divided into 16 subgroups defined by bases at positions -9 and -8 relative to designed +4 TSS, and then their TSS efficiencies were plotted. Lines represent median values of subgroups. **(F)** An A at position -8 results in different sequence preferences at position -7. The dataset of designed +1 TSSs deriving from “AYR” and “BYR” libraries was used to detect -8/-7 interaction. Calculations same as -9/-8 interaction described in **E**.

**Figure S3.**
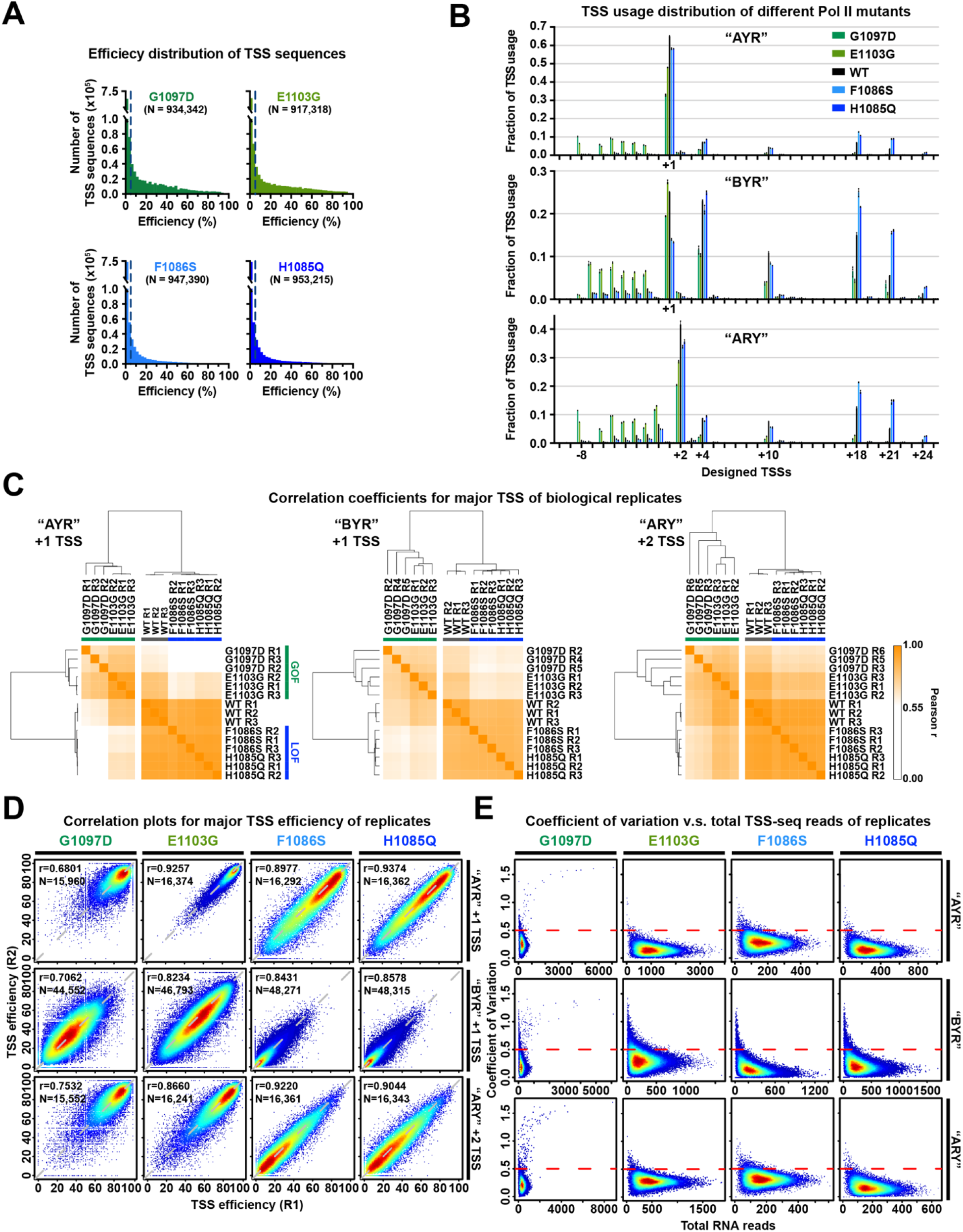
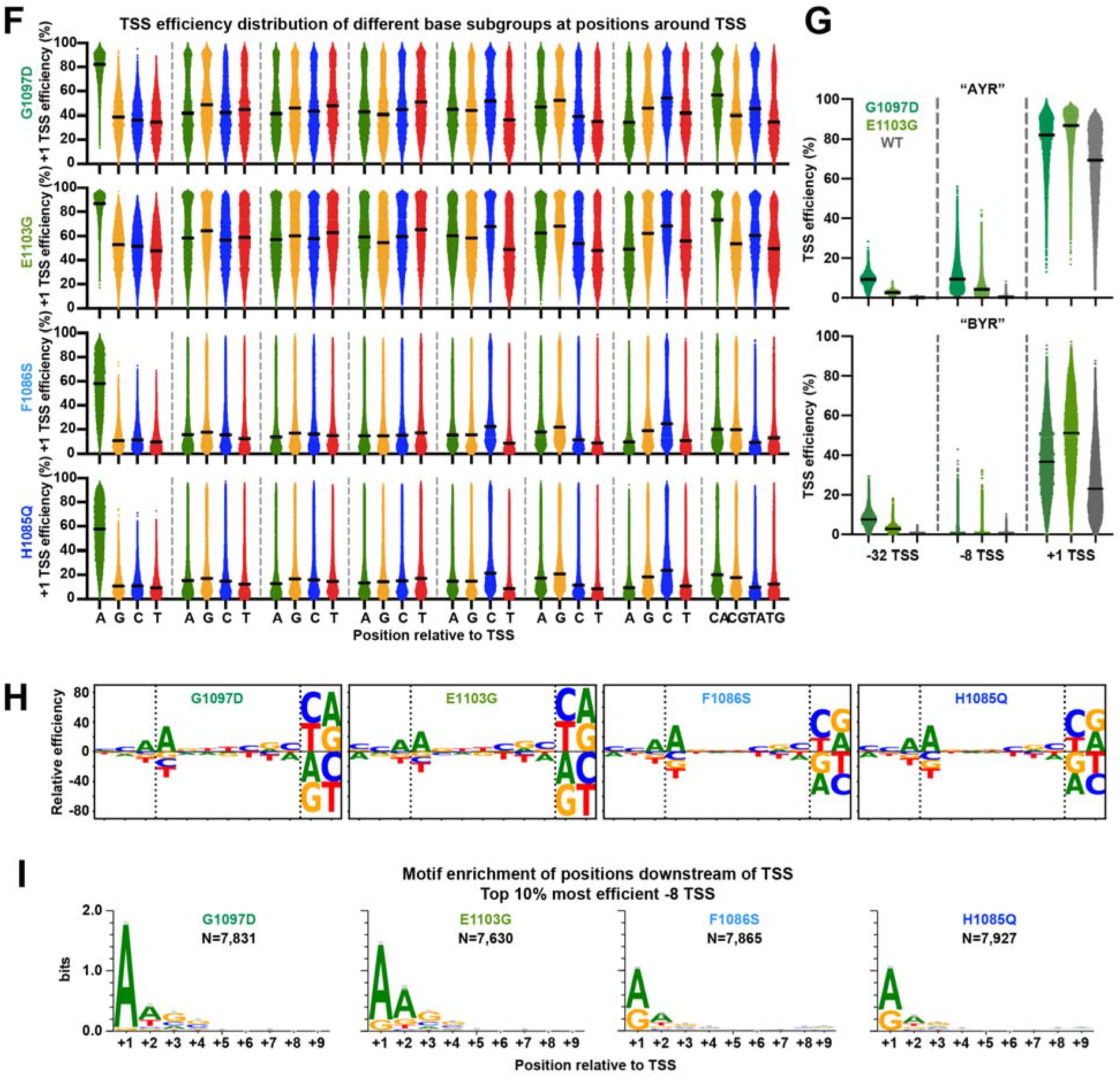
**(A)** Histograms showing the distribution of measured efficiencies for all designed -8 to +4 TSSs of all promoter variants deriving from “AYR”, “BYR” and “ARY” libraries in Pol II mutants. Dashed lines mark the 5% efficiency cutoff used to designate a TSS as active. Total numbers of TSS sequences are shown. **(B)** TSS usage distributions at designed -10 to +25 TSSs in different promoter variant “AYR”, “BYR”, and “ARY” libraries. Pol II GOF and LOF mutants shifted TSS usage upstream or downstream relative to WT, respectively. Dots represent three biological replicates. Bars are mean +/-standard deviation of the mean. **(C)** Hierarchical clustering of Pearson correlation coefficients of TSS efficiencies for the major TSSs (designed +1 TSS for “AYR” and “BYR” libraries, +2 TSS for “ARY” library) for three biological replicates for WT or mutant Pol II illustrated as a heat map. **(D)** Example correlation plots of TSS efficiency of major TSSs of promoter variants between representative biological replicates. Pearson r and number of compared variants are shown. **(E)** Plots of the coefficient of variation (CV) versus the total RNA reads for three yeast replicates in Pol II mutants. The red dashed lines mark the CV=0.5 cutoff, which was chosen as an arbitrary cutoff for variants showing reasonable reproducibility across three biological replicates. G1097D replicates contain outliers because these slow growing strains are susceptible to genetic suppressors. These outliers are filtered out using our CV cutoff because of high CVs caused by suppressor only existing in one of three biological replicates. **(F)** TSS efficiency distributions of designed +1 TSSs of Pol II mutants for base subgroups at individual positions relative to +1. Identical analysis as in **Figure S2B** for WT was performed for Pol II mutant libraries. **(G)** Pol II GOF G1097D showed greater increase in efficiency than GOF allele E1103G at upstream TSSs (designed -32 and -8 TSSs), while E1103G showed stronger effects at designed +1 TSS than G1097D. This may indicate that where Pol II is in the scanning process may affect efficiency. **(H)** Pol II initiation sequence preference in Pol II mutants. Identical analysis as in Figure 2E for WT was performed for Pol II mutant libraries. Sequence logos were generated using multiple datasets and the dashed lines indicate divisions in datasets used to generate them. Specifically, preferences at positions -11 to -9 were generated using datasets of designed +4 TSS deriving from “AYR”, “BYR” and “ARY” libraries. Preferences at positions -8 to -2 were generated using datasets of designed +1 TSS deriving from “AYR” and “BYR” libraries. Preferences at positions -1 and +1 were generated using datasets of designed +1 TSS deriving from “AYR” and “ARY” libraries. **(I)** Motif enrichment for top the 10% most efficient -8 TSSs for Pol II mutants. Identical motif enrichment analysis as in **Figure S2D** top panel for WT was performed to Pol II mutant libraries. Numbers (N) of variants assessed are indicated. Bars represent an approximate Bayesian 95% confidence interval.

**Figure S4.**
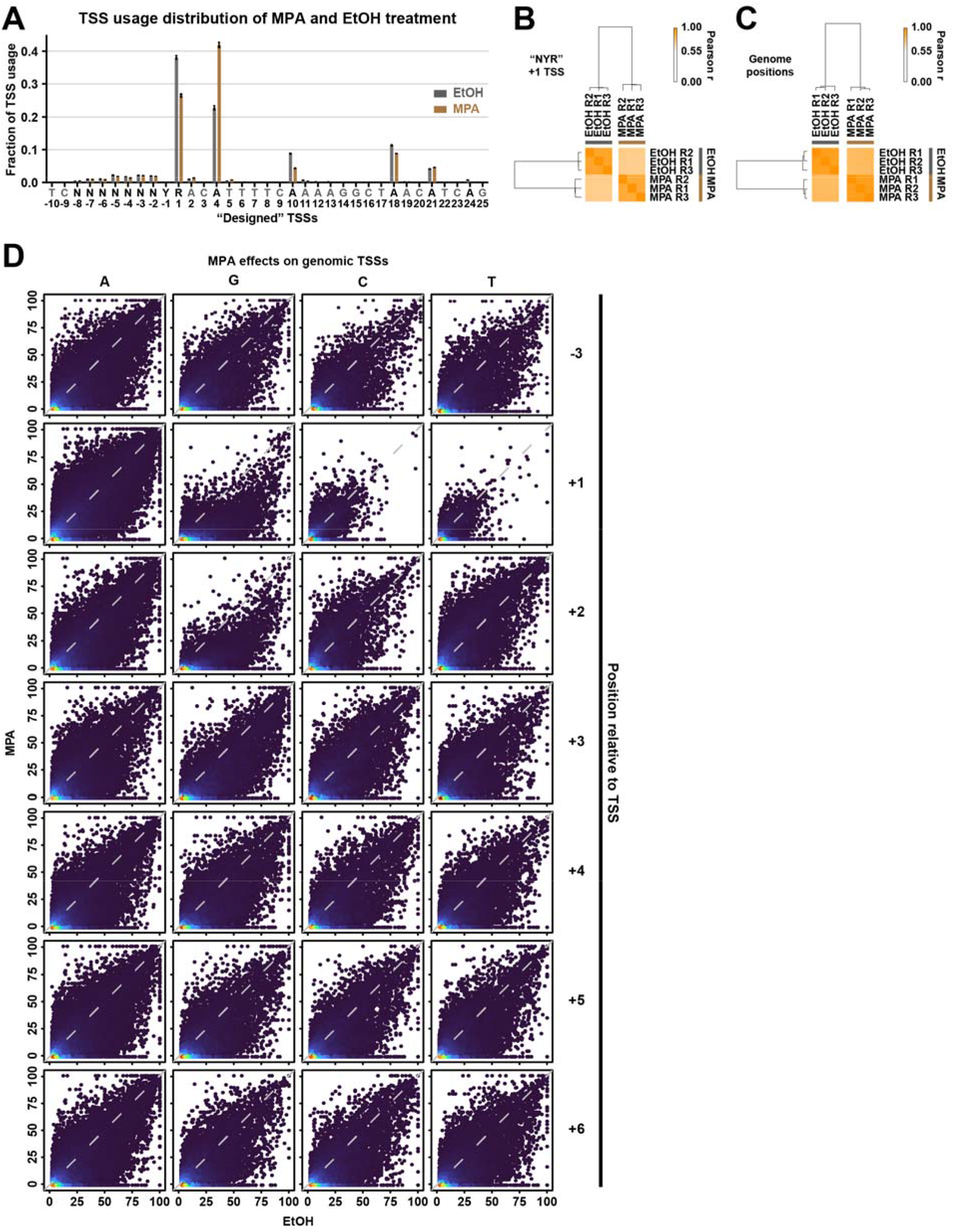
High of reproducibility of TSS usage and efficiency upon MPA treatment. **(A)** TSS usage distributions at designed -10 to +25 TSSs in WT “NYR” library (mixed AYR and BYR libraries) treated with 100% ethanol or with 20 ug/ml MPA. MPA treatment shifted TSS usage downstream relative to EtOH treatment. Dots represent three biological replicates. Bars are mean +/-standard deviation of the mean. **(B)** Hierarchical clustering of Pearson correlation coefficients of TSS efficiencies for designed +1 TSS for three biological replicates for MPA or EtOH treatment, illustrated as a heat map. **(C)** Hierarchical clustering of Pearson correlation coefficients of TSS efficiencies for all genome positions within defined promoter windows with >=3 reads in each replicate, illustrated as a heat map. **(D)** Correlation plots for combined biological replicates for TSS efficiency upon MPA treatment (*y* axes) versus EtOH treatment (*x* axes) for all TSSs ≥2% efficiency in the 25%-75% of the distribution for a curated set of 5979 yeast promoters (see Methods). TSSs are separated into groups depending on base identity at positions -3 (control) or positions +1 to +6.

**Figure S5.**
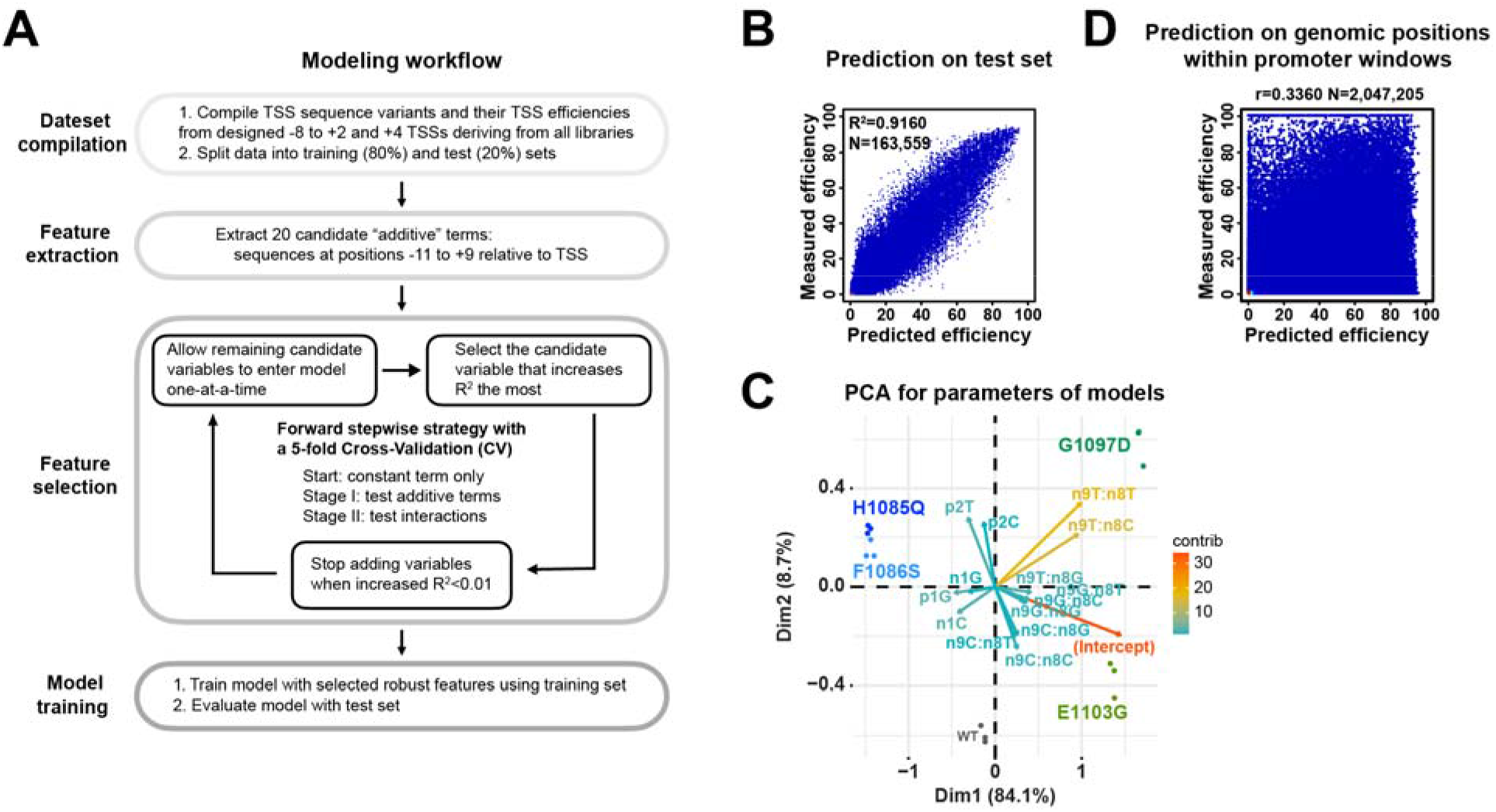
**(A)** Overview of the TSS efficiency modeling process. (1) Variants including designed -8 to +2 and +4 TSSs deriving from “AYR”, “BYR” and “ARY” libraries with available TSS efficiency were pooled for modeling. (2) Sequences at positions -11 to +9 relative to TSS of each variant were extracted. (3) To identify robust features, a forward stepwise selection strategy coupled with a 5-fold cross-validation for logistic regression was used. Data were randomly split into training (80%) and test (20%) sets. The training set was used for a stepwise regression approach that starts from a model with a constant term only and adds variables that improve the model the most one at a time, until a stopping criterion is met. In stage I, additive terms (sequences at positions -11 to +9) were tested. In stage II, interactions between positions selected in stage I were tested. Model performance was evaluated with R^2^. The stopping criterion for adding additional variables was an increase R^2^ < 0.01. (4) A logistic regression model containing selected robust features was trained with training set and then evaluated with the test set. **(B)** A scatterplot of comparison of measured efficiencies and predicted efficiencies within the test sets. Model performance R^2^ on entire test set and number of data points shown in plot are shown. **(C)** PCA analysis for parameters of models trained by the individual replicates of WT and Pol II mutant. Parameters for replicates cluster more closely than parameters for different Pol II mutants, indicating that modeling captured features of different Pol II groups and models are not overfit. The top 15 contributing variables are shown. GOF and LOF mutants were separated from WT by the 1^st^ principal component. GOF G1097D and E1103G were further distinguished by 2^nd^ principal component by additional position +2 information, which is consistent with results in **Figure S3I**, where G1097D and E1103G differentially altered +2 sequence enrichment. **(D)** A scatterplot of comparison of measured and predicted TSS efficiencies of all positions within 5979 known genomic promoter windows ^21^ with available measured efficiency. Pearson r and number (N) of compared variants are shown. Most promoter positions (82%, 1,678,406 out of 2,047,205) showed no observed efficiency, which is expected because TSSs need to be specified by a core promoter and scanning occurs over some distance downstream.

